# Genetically Corrected *RAG2*-SCID Human Hematopoietic Stem Cells Restore V(D)J-Recombinase and Rescue Lymphoid Deficiency

**DOI:** 10.1101/2022.07.12.499831

**Authors:** Mara Pavel-Dinu, Cameron L. Gardner, Yusuke Nakauchi, Tomoki Kawai, Ottavia M. Delmonte, Boaz Palterer, Marita Bosticardo, Francesca Pala, Sebastien Viel, Harry L. Malech, Hana Y. Ghanim, Nicole M. Bode, Gavin L. Kurgan, Christopher A. Vakulskas, Adam Sheikali, Sherah T. Menezes, Jade Chrobok, Elaine M. Hernández González, Ravindra Majeti, Luigi D. Notarangelo, Matthew H. Porteus

## Abstract

Recombination-activating genes (*RAG1* and *RAG2*) are critical in lymphoid cell development and function for initiating the V(D)J-recombination process to generate polyclonal lymphocytes with broad antigen-specificity. Clinical manifestations of defective *RAG1/2* genes range from immune dysregulation to severe combined immunodeficiencies (SCID), causing life-threatening infections and death early in life in the absence of hematopoietic cell transplantation (HCT). Haploidentical HCT without myeloablative conditioning carries a high risk of graft failure and incomplete immune reconstitution. The RAG complex is only expressed during the G0-G1 phases of the cell cycle at the early stages of T and B cell development, underscoring that a direct gene correction would capture the precise temporal expression of the endogenous gene, is a promising therapeutic approach for *RAG1/2*-deficiencies. Here, we report a feasibility study using the CRISPR/Cas9-based “universal gene-correction” approach for the *RAG2* locus in human hematopoietic stem/progenitor cells (HSPCs) in healthy donors and one *RAG2*-SCID patient. V(D)J recombinase activity was restored following gene correction of *RAG2*-SCID-derived HSPCs, resulting in the development of TCR αβ and γδ CD3^+^ cells and single-positive CD4^+^ and CD8^+^ lymphocytes. TCR repertoire analysis indicated a normal distribution of the CDR3 length and preserved usage of distal *TRAV* genes. We confirmed *in vivo* rescue of B-cell development, with normal IgM surface expression and a significant decrease in CD56^bright^ NK cells. Together, we provide specificity, toxicity, and efficacy data supporting the development of a gene-correction therapy to benefit all *RAG2*-deficient patients.

**KEY POINTS:**

- Human hematopoietic stem cells can be corrected to restore endogenous *RAG2* gene expression while preserving durable engraftment potential.
- Gene-corrected *RAG2* locus restores V(D)J recombination in *RAG2*-SCID patient stem cells, promoting T and B-cells’ receptor formation.

## INTRODUCTION

Severe Combined Immunodeficiencies (SCID) are genetic diseases caused by inherited mutations in genes required for T-, B-, and occasionally natural killer (NK)-cell development and function^1^. Defects in the Recombination Activating Genes 1 and 2 (*RAG1/2*) are the second most common cause of SCID. *RAG1* and *RAG2* form a heterotetrameric complex (with two copies of each protein) that initiates the V(D)J recombination process: it mediates DNA cleavage at recombination signal sequences (RSS) flanking the Variable (*V*), Diversity (*D*), and Joining (*J*) coding elements of the T-cell receptor (TCR) and immunoglobulin (Ig) receptor loci. The non-homologous end-joining (NHEJ)-mediated DNA repair of the RAG-induced DNA breaks joins the coding elements that generate functional TCR and Ig molecules.

Over 200 unique pathogenic mutations have been identified to disrupt the *RAG1/2* genes causing T^-^ B^-^ NK^+^ SCID phenotype^2,3^ and a spectrum of immune dysregulation^4,5^: Omenn Syndrome (OS), atypical SCID (AS)^6, 7^ and delayed-onset combined immunodeficiency with granulomas and/or autoimmunity (CID-G/AI)^8-10^. Patients with severe forms of *RAG*-deficiency, including SCID, OS, and AS, achieve immune reconstitution with high survival rates (>90%) following allogeneic hematopoietic cell transplantation (allo-HCT) from HLA-matched donors^11-14^. In contrast, patients who rely on haploidentical HCT without myeloablative conditioning experience high graft failure rates (>75%) or poor lymphoid reconstitution when sustained engraftment is achieved^11,12,15^. This is due to the host’s genetically defective common lymphoid progenitors (CLPs) occupying the bone marrow and thymus niches^16^ and preventing donor’s CLPs from establishing lymphopoiesis.

To identify therapeutic options beneficial to all *RAG*-deficient patients, feasibility studies using lentivirus-based gene therapy were performed in *Rag1/2* disease mouse models^17-19^. These studies showed variable therapeutic outcomes suggesting that overexpressing *Rag* genes could not robustly and faithfully recapitulate the levels of regulation and expression necessary to support immune system development and function, thus presenting significant concerns for clinical translation.

Advances in gene-editing technology^20-22^ have allowed gene-correction of human HSPCs. This strategy offers significant therapeutic benefits for diseases like *RAG1/2*-deficiency, where the underlying gene relies on strict spatiotemporal gene regulation and expression^23^. The RAG protein complex is only expressed in G_0_/G_1_ in early T and B-cell development, thus demonstrating precise cell cycle and developmental specificity. Cluster Regulatory Interspaced Short Palindromic Repeats-associated Cas9 nuclease (CRISPR/Cas9) is a genome engineering platform^21-23^ that has been successfully applied to primary human cells^24-26^, including HSPCs^26-30^ to introduce a desired genomic modification through homologous recombination-mediated gene-targeting (HR-GT)^23^. We have previously demonstrated *in vitro* rescue of the T-cell developmental blockage using CRISPR/Cas9-AAV6 based correction of a patient’s *RAG2* homozygous null mutation (c.831T>A; p.Y277*)^31^. However, the *in vivo* efficacy of the gene-correction approach remains to be determined. Here, we develop and test the safety and effectiveness of a novel approach for correcting *RAG2*-deficiency in a patient’s own HSPCs. We used the CRISPR gene-correction platform to repair the DNA sequence of a *RAG2*-deficient patient’s stem cells carrying two null compound heterozygous variants causing *RAG2*-SCID and show that it can restore V(D)J activity, rescue T-and B-cell development, and correct the immature NK cell phenotype *in vivo*, following long-term engraftment into an immunodeficient NSG-SGM3 mouse model.

## METHODS

### Human CD34^+^ HSPCs

PB-CD34^+^ cells from a *RAG2*-SCID subject were collected following informed consent (protocol NCT00001405, approved by the NIH IRB as NIH protocol 94-I-0073). Additional information is detailed in supplemental methods.

### Off-target activity assay

The top forty-eight putative off-target sites for *RAG2* were identified using COSMID^32^ and tested using rhAmpSeq^33,34^ (Integrated DNA Technologies). Additional information is described in supplemental methods.

### Colony-forming units (CFU) assay

Single live HSPCs were sorted and analyzed, as previously described^27^. Colony genotyping is detailed in supplemental methods.

### Transplantation of human gene-modified CD34^+^ HSPCs and engraftment assessment

Rescue experiments, detailed in supplemental methods, were done using frozen PB-HSPCs from one *RAG2*-SCID patient with compound heterozygous null variants (c.296C>A, c.1324C>A; p.P99Q, A442T).

### T and B-cell receptors repertoire analysis

High-throughput sequencing of TCRα and TCRβ and immunoglobulin M (IgM) heavy chain (Vh) analysis was performed as described in the supplemental methods. PCR assay for the seven Vh chains was performed as described^35^.

### Karyotype analysis

*RAG2*-SCID patient-derived PB-HSPCs mock or RNP only treated were subjected to G-band karyotyping analysis on 20 cells derived from each condition [WiCell Cytogenetics (Madison, WI, USA)].

### Statistical analysis

Statistical analysis was done with Prism 9 (GraphPad software).

### Ethics and animal approval statement

All the work described in this study was carried out in compliance with all relevant ethical regulations. The animal studies were reviewed, approved, and monitored by the Stanford University IACUC committee.

### Data sharing statement

Off-target sequencing data have been deposited at the BioProject (ID PRJNA817273) under accession code SAMN26753110 (https://www.ncbi.nlm.nih.gov/biosample/26753110). The authors declare that the data supporting this study are available within the paper and its supplementary information files or from the authors upon reasonable request.

## RESULTS

### Efficient “universal” gene-targeting strategy of *RAG2* to the endogenous locus in human HSPCs

Two central concepts of the corrective therapeutic approach for *RAG2*-deficiency are that hematopoietic stem cells (HSCs) and their progeny will preserve the physiological gene regulation necessary to achieve temporal and lineage-specific RAG2 protein expression and activity (Figure 1A-B) and that most of the *RAG2*-causing mutations, including deletions, will be corrected. We screened and selected a *RAG2*-specific guide (sgRNA-3, supplemental Figure 1A-B) that when complexed with HiFi Cas9 nuclease achieves a median INDEL frequency (gene-editing mediated NHEJ; GE-NHEJ) of 52.6% (range: 27.1-91.0%) in CB-HSPCs and 68.0% (range: 31.0-81.0%) in PB-HSPCs. The guide’s INDEL profile is characterized by 2-17 bp deletions (supplemental Figure 1C, 1F), which abrogate RAG2 protein expression *in vivo* (supplemental Figure 1D-E).

**Figure 1.**
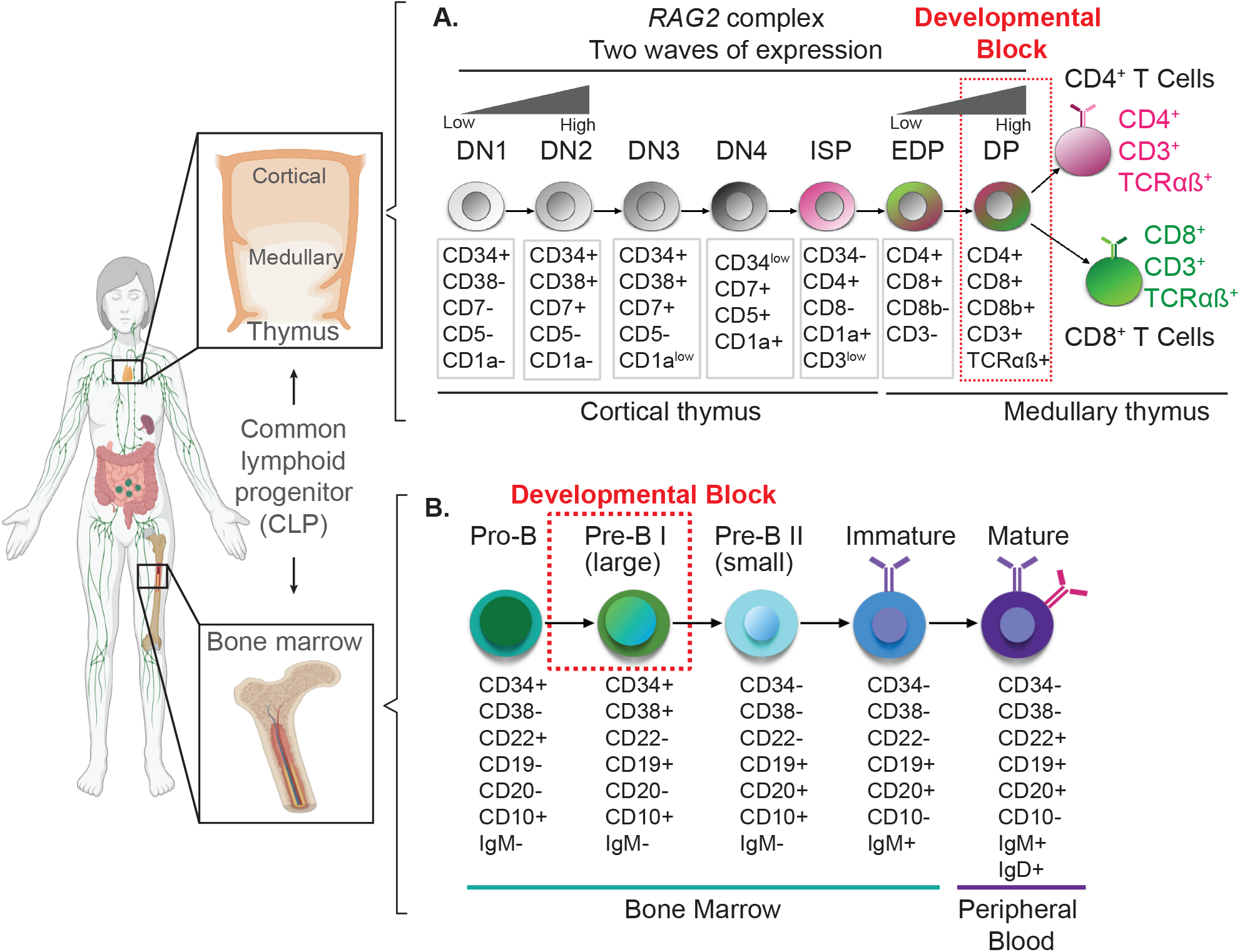
Schematics of the hematopoietic developmental defect in RAG2-SCID patients. **(A)** Overview of the human T-cell (top panel) and **(B)** B-cell (bottom panel) developmental stages in thymus and bone marrow from common lymphoid progenitor (CLP). Dotted red squares mark the developmental block in RAG2-SCID patients. RAG2, recombination activating gene 2; DN, double negative stage; ISP, immature single positive stage; EDP, early double-positive stage; DP, double-positive stage.

We used adeno associated virus 6 (AAV6) to deliver a codon-optimized (*coRAG2)* transgene bearing homology arms to the guide’s cut sites (Figure 2A). Absolute quantification showed an average of 40.6 ± 3 (s.e.m, n=13 unique HSPCs donors) alleles carrying the co*RAG2* (Figure 2B) or higher (1.4 -1.7-fold increase) when total alleles were quantified at single-cell resolution (Figure 2C, green and hashed bars combined; supplemental Figure 2A-B; supplemental Figure 3A-B): 58.0% [(5,000 MOI, n = 5 donors); 56.0% (2,500 MOI, n = 4 donors); 68.7% (1,250 MOI, n = 2 donors). We estimated the distribution of cells with one (mono-allelic) or two (bi-allelic) alleles targeted as a function of viral multiplicity of infection (MOI) and observed an increase in mono-allelic HR-GT with decreased MOI (Figure 2C): 5,000 MOI 25.6 ± 2.0 (s.e.m.) mono-allelic and 32.4 ± 8.7 (s.e.m.) bi-allelic; 2,500 MOI 31.7 ± 4.5 (s.e.m.) mono-allelic and 24.5 ± 7.4 (s.e.m.) bi-allelic; 1,250 MOI 38.6 ± 14.2 (s.e.m.) mono-allelic and 30.1 ± 4.6 (s.e.m.) bi-allelic. Importantly, correction of a single allele is sufficient to revert the disease phenotype.

**Figure 2.**
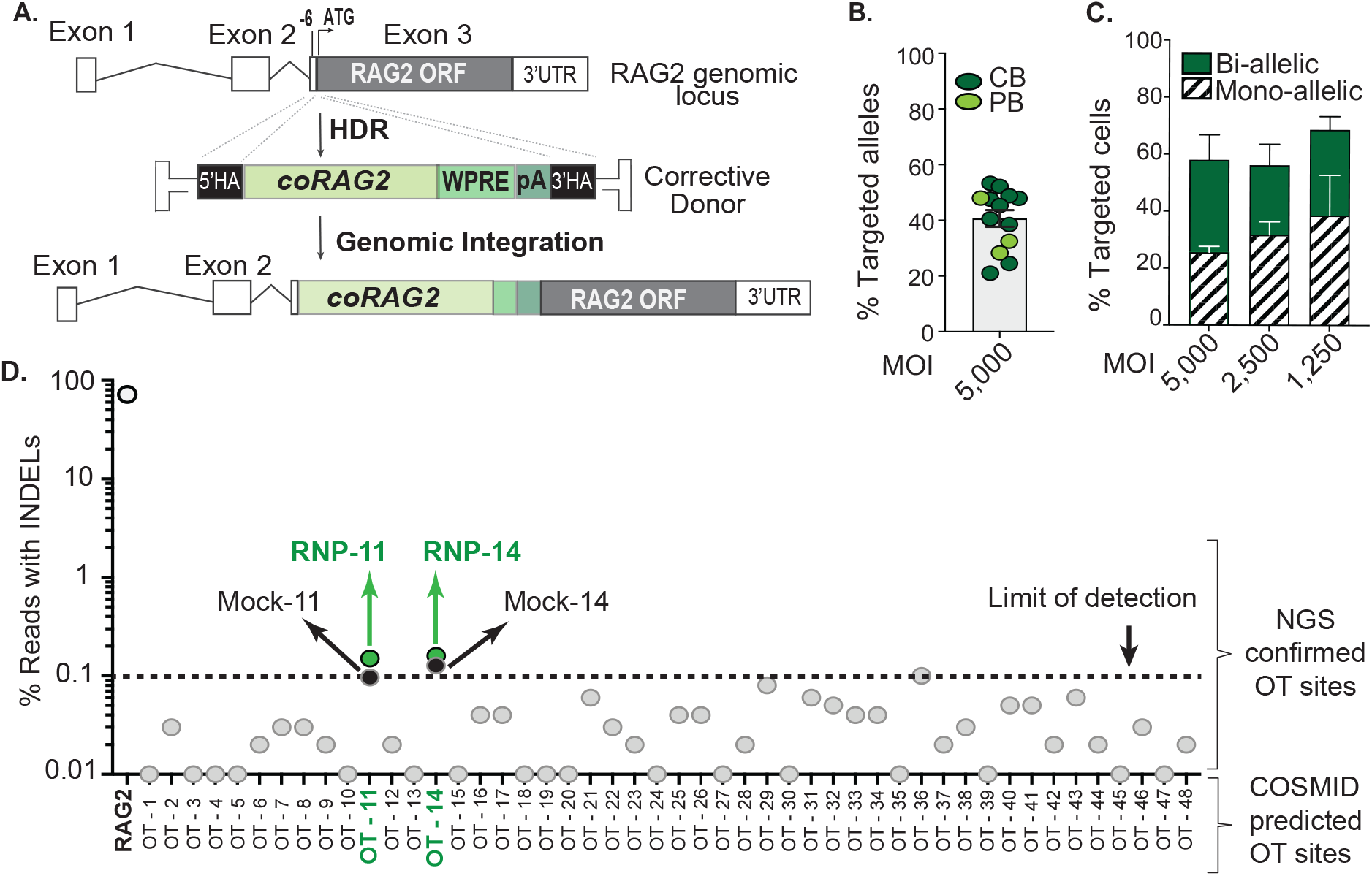
Efficacy and safety of genome targeting the RAG2 locus using “universal” correction strategy. **(A)** Schematics of gene-targeted integration of codon-optimized RAG2 cDNA (coRAG2) and expression cassette. coRAG2 sequence is under the control of the endogenous promoter. **(B)** Percent coRAG2 gene-targeted (coRAG2-GT) in healthy donor (HD)-purified hematopoietic stem cells (HSPCs) from fresh cord blood (CB) and frozen peripheral blood (PB). Each circle represents a unique human HSPC donor. Genome targeted integration was quantified by digital-droplet PCR (ddPCR). **(C)** Frequency of cells with one (mono-allelic) or two (bi-allelic) alleles targeted, as a function of virus’ MOI. Analysis was done on single cells sorted onto methylcellulose plates (n = 5 - 5,000 MOI; n = 4 - 2,500 MOI; n = 2 - 1,250 MOI). Bars ± s.e.m. **(D)** Off-target analysis using RAG2-SCID (c.296C>A; c.1342C>A) patient-derived HSPCs. Next-generation sequencing (NSG) of 48 COSMID predicted off-target (OT) sites in edited-only (RNP-sgRNA guide #3 and HiFi Cas9 nuclease) or electroporated-only (mock, nucleofected without RNP). Shown INDELs reads for on target (RAG2 gene, white circle), OT sites below the limit of detection (grey circles), and above the limit of detection (green circles).

We report no significant bias in the co*RAG2*-modified HSPCs’ (co*RAG2*-HSPCs) ability to generate all four early myeloid and erythroid hematopoietic progenitor cells compared to control conditions, demonstrating an absence of unintended perturbations in the non-lymphoid lineages (supplemental Figure 2A-C). To address the virus-related toxicity at higher MOIs, we tested p53 binding protein 1 (53BP1) inhibitor (i53) in combination with a lower AAV6 concentration. Treatment with i53 inhibits the NHEJ pathway and tips the balance towards homology direct repair (HDR) pathway^36^. We observed a 1.5-fold higher HR-GT without diminishing the modified cells’ differentiation potential^37^ (supplemental Figure 2B). Overall, we show that co*RAG2-*HSPCs support early hematopoietic progenitor development.

### Specificity and safety of the *RAG2* gene-correction approach

To evaluate the off-target profile of the *RAG2* lead guide (sgRNA-3), we used COSMID^32^, a bioinformatic tool to predict the putative guide’s binding sites, and rhAMP-Seq^33,34^ to confirm them. Off-target activity at a total of 48 predicted loci was quantified^38^ in the genome of PB-derived HSPCs from a *RAG2*^null^ patient carrying compound heterozygous missense mutations (c.[296C>A; 1342C>A]) (Figure 2D). We compared the percent INDELs (GE-NHEJ) detected in cells electroporated with the ribonucleoprotein (RNP) to mock-treated (control) cells. Sites considered true off-target (OT) were those with GE-NHEJ > 0.1% in the RNP-treated condition relative to the mock sample. None of the sites were valid off-targets (Figure 2D; Table 1). While OT-11 and OT-14 are not *bona fide* off-target sites because there is no increased frequency of INDELs compared to Mock, in Table 1, we annotate these sites for completeness. These results, combined with no chromosomal abnormalities detected by karyotype analyses (supplemental Figure 4A-B), suggest a high degree of specificity by the lead *RAG2* guide (sgRNA-3) when complexed with HiFi *S*.*p*.Cas9^22^ (RNP).

**Table 1.**
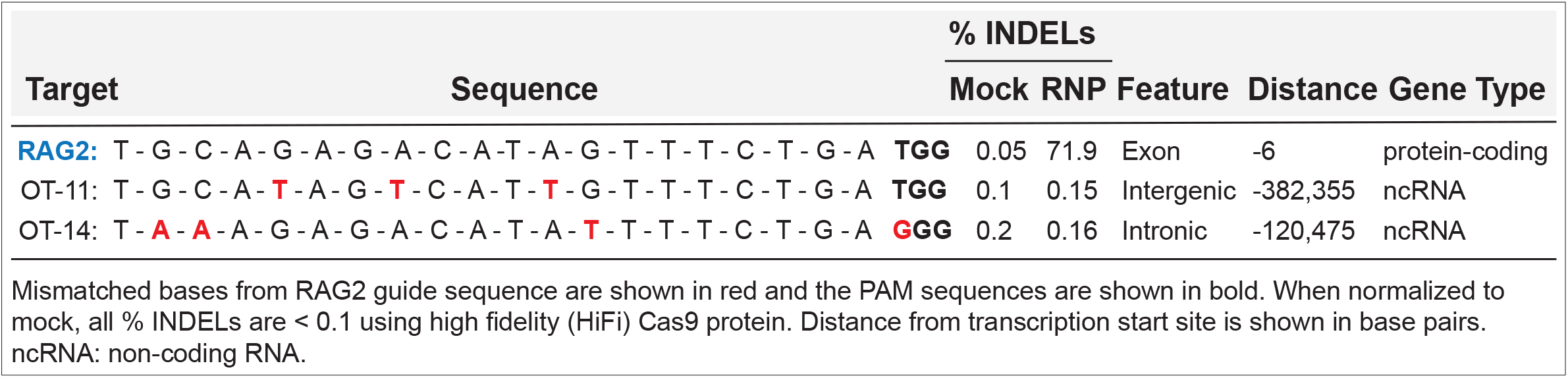
Description of off-target (OT) sites above limit of detection (>0.1)

Cumulatively, we transplanted 63 mice with 28 million gene-modified HPSCs (3.5 million modified cells derived from healthy donors and 24.5 million modified cells derived from *RAG2*-SCID patients) and analyzed them 18-22 weeks following transplantation. We observed no hematopoietic skewing or detectable gross tumors. In sum, these data offer strong genetic and functional evidence for the safety of our gene-correction strategy for *RAG2*-deficiency.

### Human hematopoietic system is derived from co*RAG2*-HSPCs

To determine if gene-modified stem cells can support long-term normal hematopoiesis, we performed xenotransplantation studies into immunodeficient NSG or NSG-SGM3 mice (Figure 3A). We first tested the hematopoietic reconstitution of co*RAG2*-HSPCs in fresh CB-HSPCs using intra-hepatic (IH) injections into NSG neonates because this system has been shown to best support lymphopoiesis^39^. Primary human engraftment was measured 18 weeks post-transplant by FACS analysis for human chimerism (% hCD45^+^ HLA A-B-C^+^ cells) in the bone marrow (Figure 3B left panel; supplemental Figure 5A) and spleen (Figure 3B right panel; supplemental Figure 5B), as follows: unmodified 28.6% (range: 10.3-39.4%), RNP 15.4% (range: 4.4-41.8%), 15.8% (range: 7.0-40.3%). We observed no difference in the repopulation capacity of HSPCs that underwent HR-GT or GE-NHEJ compared to cells that did not (mock or wild type). Within the hCD45^+^HLA^+^ cells, the mean frequency of cells expressing co*RAG2* purified from bone marrow (BM) was 19.5 ± 6.5 (s.e.m.; n = 10) and from spleen was 20.7 ± 5.3 (s.e.m, n = 11), as compared to 21.0% bulk targeted alleles before engraftment (Figure 3D -Tx vs +Tx). co*RAG2* cDNA was also detected in sorted lymphocytes (CD19^+^ B cells and CD3^+^ T cells) from BM (Figure 3D, yellow squares and triangles) and spleen (Figure 3D, red squares and triangles). The quantification only showed a proliferative advantage in T-cells sorted from spleen [53.5 ± 6.1 (s.e.m., n = 4)] with the remaining sorted cells expressing the transgene at levels comparable to bulk (not sorted) human hCD45^+^HLA^+^ cells: 16.6 (n = 1; sorted CD19^+^ from BM); 27.9 ± 9.9 (s.e.m., n = 3 sorted CD19^+^ from spleen); 53.6 ± 6.2 (s.e.m., n = 4; sorted CD3^+^ from BM); 34.4 ± 11.5 (s.e.m., n = 5; sorted CD3^+^ from spleen), (Figure 3D).

**Figure 3.**
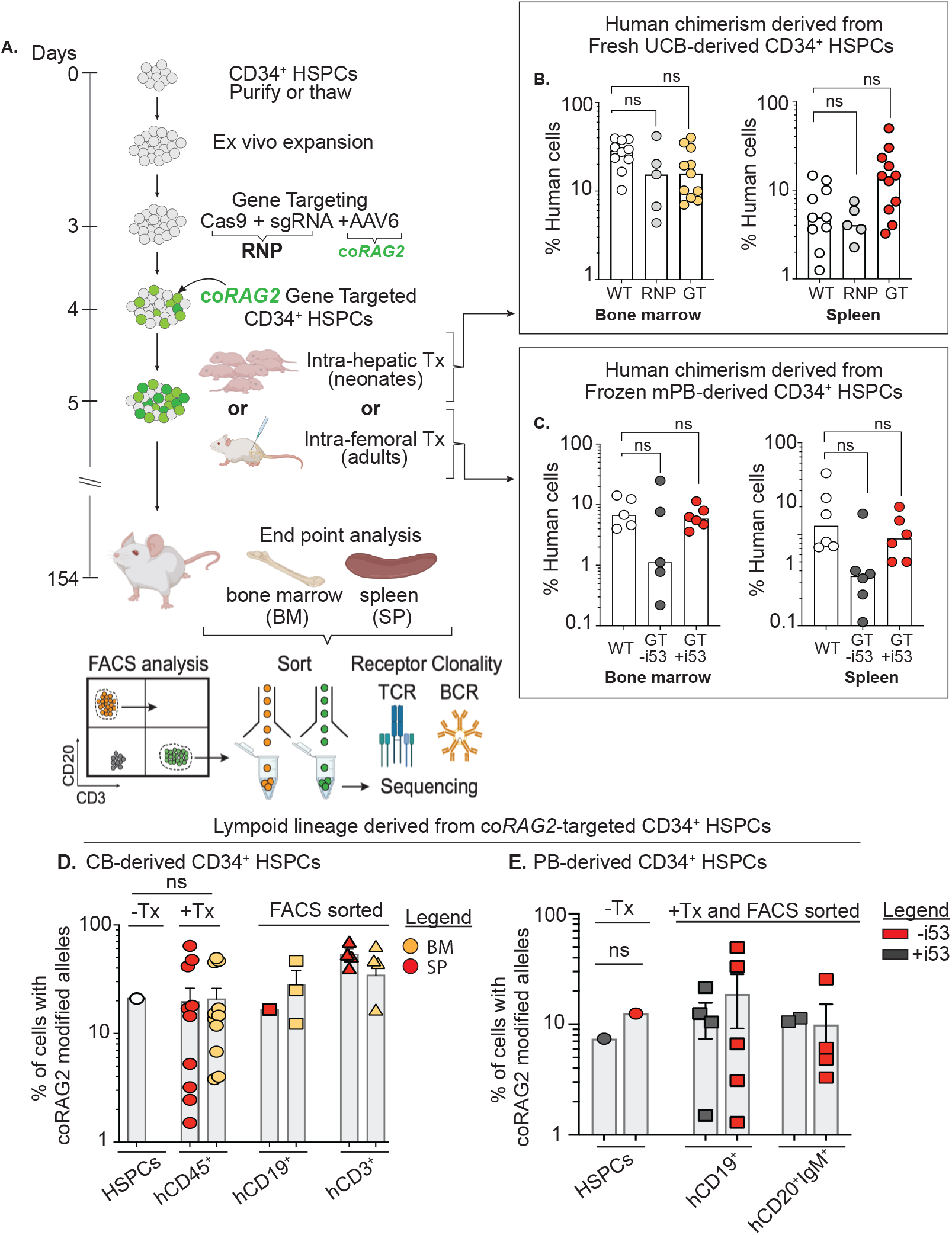
In vivo lymphoid lineage development from coRAG2-targeted healthy donor-derived HSPCs. **(A)** Schematic of engraftment protocol of human HSPCs into NSG immunodeficient mice and secondary analysis. **(B)** Percent human cell chimerism (CD45^+^ HLA A-B-C^+^ double-positive cells) in bone marrow (BM) and spleen (SP) of mice 22-weeks post-transplant with CB (intra-hepatic injection) or **(C)** PB-derived HSPCs (intra-femoral injection) edited with RAG2-sgRNA guide 3 (RNP, light grey circles in panel B) or targeted with coRAG2 cassette (GT, yellow and red circles in panel B, dark grey and red circles in panel C). Each dot represents an individual mouse; Panel B: WT (n=10), RNP (n=5), GT (n=11); Panel C: WT (n=5), GT-i53 (n=5), GT+i53 (n=6). Bars = median **(D)** Percent coRAG2-GT by ddPCR before (-Tx) and after (+Tx) engraftment of CB and **(E)** PB-derived CD34^+^ HSPCs into NSG mice and FACS sort, as marked. Bars: mean ± s.e.m.; -i53, without p53 inhibitor; +i53, with p53 inhibitor; stats. One-Way ANOVA, non-parametric test, Kruskal-Wallis test, Dunn’s multiple comparisons test.

To assess the efficacy of the gene-correction strategy in a cell source of similar derivation as that of *RAG2*-deficient patients, we repeated the xenotransplantation studies using frozen PB-HSPCs derived from healthy donors. An equal number of unmodified (wild type and mock) and HR-GT PB-HSPCs treated (+i53) or not (-i53) were transplanted intra-femoral into sub-lethally irradiated 8 weeks-old NSG-SGM3 mice (Figure 3A, 3C). Twenty-two weeks post-transplantation, the median frequencies of hCD45^+^HLA^+^ cells in BM were as follows: unmodified 6.8% (range: 4.0-14.2); HR-GT (-i53) 1.12% (range:0.2-25.0); HR-GT (+i53) 5.9% (range: 3.6-11.4). We observed no statistical difference in the repopulation capacity of cells that underwent HR-GT compared to cells that did not, nor among HR-GT cells treated or not with p53 inhibitor (i53) in BM (Figure 3C left panel; supplemental Figure 6A-B) or SP (Figure 3C right panel). Human cells carrying the co*RAG2* cDNA were detected among sorted mature B cells: CD19^+^ [-i53: 11.5 ± 4.1 (s.e.m., n = 4); +i53: 18.8 ± 9.6 (s.e.m., n = 5)] and CD20^+^IgM^+^ [-i53: 10.9 ± 0.4 (s.e.m., n = 2); +i53: 9.9 ± 5.2 (s.e.m., n = 4)], providing the first line of evidence that co*RAG2* cDNA drives productive V(D)J recombination reactions (Figure 3E).

To assess the HR-GT cells’ ability to differentiate into multiple hematopoietic lineages, we used human CD3, CD19, CD14, and CD235a markers to identify lymphoid (B, T), myeloid (monocytes), and erythroid population, respectively. Compared to unmodified (mock) cells, co*RAG2-*modified cells showed no statistical difference skewing towards a particular lineage in BM (supplemental Figure 5C left panel), in circulating cells in the peripheral blood (supplemental Figure 5C right panel) or spleen (supplemental Figure 5D left panel; supplemental Fig. 7). Together, these data demonstrate that the co*RAG2* transgene can support durable hematopoiesis and drive lymphoid development, *in vivo*, without lineage skewing.

### In vivo B-cell lineage reconstitution from gene-corrected *RAG2*^null^ HSPCs

We performed xenotransplantation experiments into NSG-SGM3 immunodeficient mice to assess the phenotypic correction of the B-cell developmental defect in *RAG2*^null^ patient HSPCs (*RAG2*^null^ HSPCs). Four different non-GMP (good manufacturing practice) grade AAV6 virus batches carrying the co*RAG2* transgene (AAV6^co*RAG2*^) were purified and tested (5,000 MOI) on frozen *RAG2*^null^ HSPCs (Figure 4A). Total HR-GT (green bars) or GE-NHEJ (grey bars) generated by three different virus batches in the absence of i53 treatment showed a mean % INDEL of 54.9 ± 3.7 (s.e.m.), mean %HR of 29.9 ± 5.1 (s.e.m.), respectively, with 15.2% alleles (± 5.6 s.e.m.), remaining unmodified. These viruses, however, decreased cellular viability, diminished CFU, and long-term engraftment potential of gene-corrected *RAG2*^null^ HSPCs (data not shown). We purified a fourth AAV6^co*RAG2*^ batch and used it to transduce the patient cells at lower MOI (2,500) and in the presence of p53 inhibitor (+i53). Under these optimized conditions, we achieved a 19.2% HR-GT and 37.0% GE-NHEJ with no observed toxicity.

**Figure 4.**
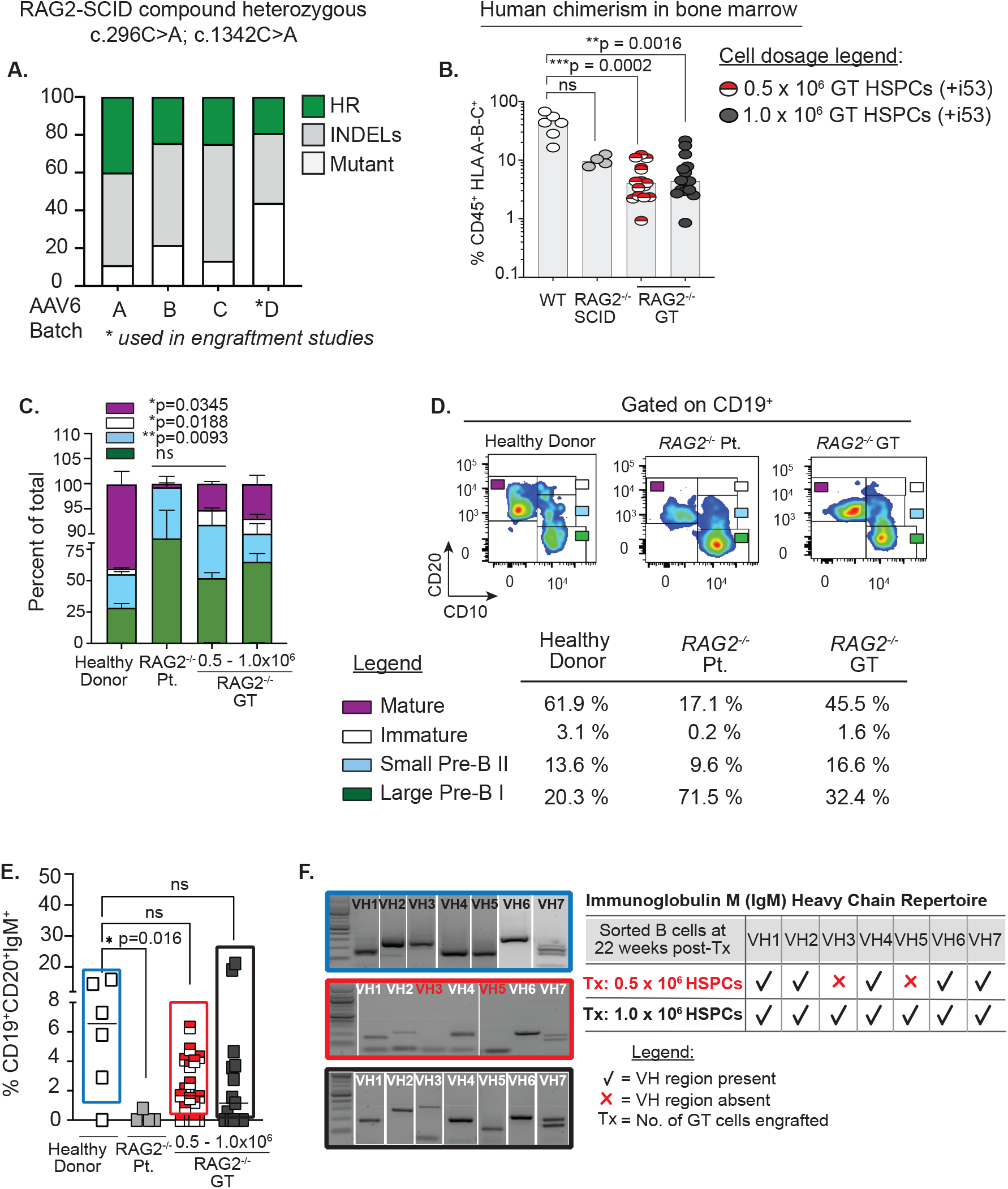
Ex vivo gene-targeted RAG2^null^ HSPCs correct in vivo B-cell developmental block. **(A)** Percent of total genome editing (INDELs and HR) in RAG2^null^ patient-derived HSPCs, using four different AAV6 production lots. AAV6 lot D (asterisk marked) was used for all subsequent engraftment studies. **(B)** Percent human cells (CD45^+^ HLA A-B-C^+^) engrafted in BM after transplanting (22 weeks post-Tx) 0.5 ×10^6^ un-corrected RAG2^null^ patient-derived HSPCs (n=4 mice, grey circles), 0.5 ×10^6^ coRAG2-GT HSPCs (n=15 mice, half red circles) or 1.0 ×10^6^ coRAG2-GT HSPCs (n=15 mice, black circles). Healthy donor (HD) HSPCs were used as control (n=6, white circles). RAG2^null^ patient genotype: c.296C>A; c.1342C>A. **(C)** FACS-based quantification is shown in **(D)** of large Pre-B I, small Pre-B II, immature, and mature B cells derived from coRAG2-GT HSPCs. Each population is graphed as a percent of total B cells. Bars: mean ± s.e.m. **(D)** Representative FACS plots of B-cell developmental stages derived from a healthy donor (left panel), RAG2^null^ patient (middle panel), and coRAG2-GT RAG2^null^ HSPCs. FACS-based quantification of percent cells in each developmental stage is shown. **(E)** FACS-based quantification of CD19^+^CD20^+^IgM^+^ triple-positive B cells derived from each condition tested. **(F)** PCR-based sequencing of immunoglobulin M (IgM) heavy chain (Vh) families from sorted triple-positive B-cells.

Two different cell doses (5.0 × 10^5^ and 1.0 × 10^6^) of HR-GT *RAG2*^null^ HSPCs were transplanted intra-femoral into NSG-SGM3 mice and unmodified controls (mutant and wild type cells). Human chimerism (hCD45^+^HLA^+^) was quantified at 22-weeks post-transplant in the BM and resulted as follows: 4.4% (range: 0.9-21.7; Tx = 1.0 × 10^6^ HR-GT) in mice transplanted (Tx) with 1.0 × 10^6^ HR-GT; 4.1% (range: 0.9-12.3; Tx = 5.0 × 10^5^ HR-GT); 9.6% (range: 7.78-12.5; Tx = 1.0 × 10^6^ *RAG2*^null^ HSPCs); and 43.5% (range: 16.4-67.3; 1.0 × 10^6^ healthy donor HSPCs) (Figure 4B). A two-fold decrease in human chimerism was observed between the healthy donor and HR-GT *RAG2*^null^ HSPCs, regardless of the cell dose transplanted. We observed no difference in the level of engraftment between the healthy donor and unmodified *RAG2*^null^ HSPCs (Figure 4B white and grey circles), suggesting an increased sensitivity of *RAG2*^null^ HSPCs to the gene-correction reagents. Transplanting NSG-SGM3 mice with a lower dose of HR-GT *RAG2*^null^ HSPCs generated a statistically significant increase in the fractions of small pre-B II (light blue bar) cells, immature (white bar), and mature (burgundy bar) cells when compared to uncorrected mutant cells (Figure 4C-D). The triple-positive CD19^+^CD20^+^IgM^+^ mature B cells derived from HR-GT *RAG2*^null^ HSPCs (Figure 4E, red/white and black squares) developed at comparable levels to healthy donor engrafted cells (Figure 4E, white squares). When sorted (supplemental Figure 8) and analyzed for IgM immunoglobulin heavy chain repertoire, the HR-GT *RAG2*^null^ HSPCs engrafted with a higher cell dose (1.0 × 10^6^) showed expression of all 7-immunoglobulin heavy chain Variable gene families (Figure 4F, black bottom panel). The lower cell dose (5.0 × 10_5_) expressed 5 out of 7 heavy chain variable regions in the (Figure 4F, middle red panel). Together, this data shows that up to 84.8% of *RAG2*^null^ alleles are gene-modified, of which up to 29.9% alleles are corrected and up to 54.9% alleles are disrupted. Furthermore, HR-GT *RAG2*^null^ HPSCs can restore the B-cell development block.

### Transplantation of *RAG2*^null^ gene-corrected HSPCs rescues T-cell development in NSG-SGM3 mice

Since we observed correction in the B-cell compartment, we also assessed the potential of the HR-GT *RAG2*^null^ HSPCs in rescuing the T-cell developmental defect. Despite lower human chimerism in the spleen (Figure 4B, 5A), corrected *RAG2*^null^ HSPCs gave rise to CD3^+^ T-cells at levels comparable to healthy donor-derived T-cells (in 3 mice) (Figure 5B-C). The derived CD3^+^ T-cells express αβ and γδ T-cell receptors (TCR) and gave rise to single-positive CD4^+^ and CD8^+^ T-cells, demonstrating that CLPs derived from corrected RAG*2*^null^ HSPCs undergo normal lymphopoiesis and achieve maturation.

**Figure 5.**
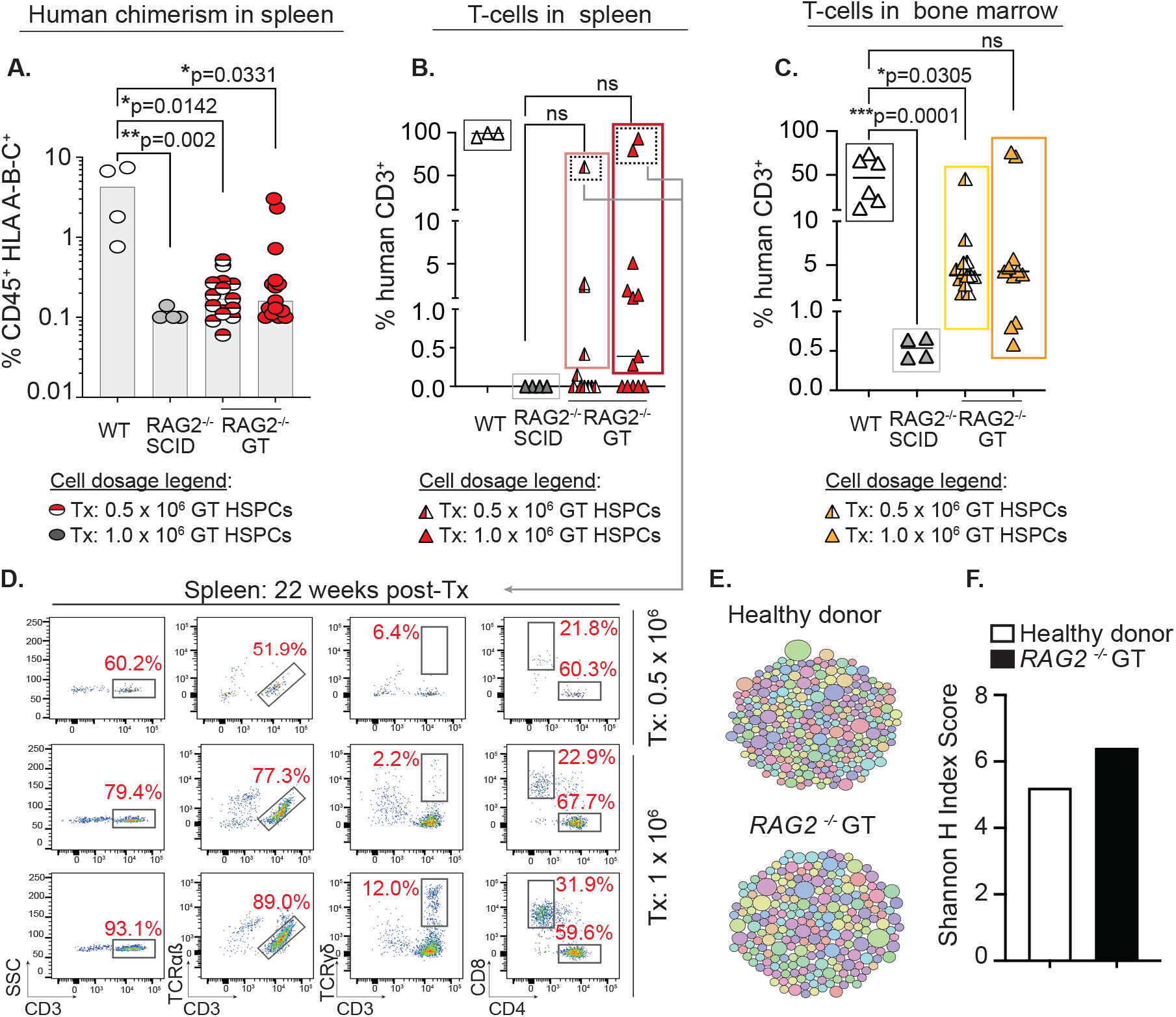
Correction of RAG2 gene function in RAG2null HSPCs restores V(D)J activity and normal T-cells development. **(A)** Percent human cells (CD45^+^ HLA A-B-C^+^) detected in spleen (SP) (22-weeks post-Tx) with coRAG2-GT RAG2^null^ HSPC (0.5 × 10^6^, half-colored red circles or 1.0 × 10^6^, full-colored red circles). HD (white circles) and un-corrected RAG^-/-^ (grey circles) HSPCs-derived human cells were engrafted and analyzed in parallel. **(B)** Human CD3^+^ T-cells detected in the spleen (SP) and **(C)** bone marrow (BM) derived from coRAG2-GT RAG2^null^ HSPCs. **(D)** FACS plots showing T-cells analysis derived from 3 mice with the highest level of human CD3^+^ cells (dotted squares). Functional V(D)J rearrangement is demonstrated by the presence of CD3^+^TCR α/β, CD3^+^TCR γ/δ, and single-positive CD4^+^ and CD8^+^ derived from coRAG2-GT RAG2^null^HSPCs. **(E)** Treemap diversity analysis for TCRA/TCRD CDR3 sequences from sorted CD3^+^ cells from (C). Each circle is a unique CDR3 sequence, and the size of the circle represents the frequency out of the total number of reads. **(F)** Shannon H index score quantification of CDR3 sequence from (E-F), showing oligoclonal repertoire. Shannon index score of ⊠8 indicates polyclonal repertoire. Stats. One-way ANOVA, nonparametric, Kruskal-Wallis test. Median plotted in A-C.

We used high-throughput sequencing (HTS) to quantify the diversity and composition of the TCR repertoire and determine the clonality at the complementary determining region 3 (CDR3) level. CDR3 is the region that promotes TCR-peptide MHC complex formation. Using a treemap profile analysis of the CDR3 of sorted HR-GT *RAG2*^null^ derived CD3^+^ T cells and healthy donor control, we determined an oligoclonal pattern of TCR CDR3 specificity, with Shannon’s H entropy index of 6.4 and 5.2, respectively (Figure 5E-F). Virtual spectratyping analysis revealed similar CDR3 lengths of productive rearrangements in bulk T-cells from HR-GT *RAG2*^null^ and healthy donor control (supplemental Figure 9).

It has been reported that mature T cells derived from *RAG2* patients carrying hypomorphic mutations have an abnormal *TRA* repertoire composition defined by a markedly reduced usage of the most 5’ *TRAV* and 3’ *TRAJ* genes^40^. HTS of TCR rearrangements at the TCRα/TCRδ (*TRA*/*TRD*) loci were demonstrated. HR-GT showed a similar rearrangement pattern of *TRAV* and *TRAJ* gene usage in *RAG2*^null^*-*derived CD3^+^ T cells and healthy donor-derived CD3^+^ T-cells (supplemental Figure 10A-B). Analysis of *TRB* rearrangements showed comparable patterns in HR-GT *RAG2*^null^ derived CD3^+^ T cells to healthy donor control (supplemental Figure 10C-D). Together, the data demonstrate that the function of the RAG complex is restored following endogenous genomic integration, promoting T-cell development with a normal VJ pairing in gene-edited cells, supporting the previously published *in vitro* results^31^.

### Correction of the immature NK CD56^bright^ phenotype in *RAG2*^null^ patient

It was reported that *RAG*-deficient mice display an increased proportion of natural killer (NK) cells that have an immature phenotype, with heightened cytotoxic activity and limited survival potential^41^. These observations were also reported in patients with SCID caused by defects in *RAG* and NHEJ genes^42^. The FACS analysis of human chimerism in mouse BM confirmed that NK cells derived from *RAG2*^null^ HSPCs expressed significantly higher CD56brightCD16-cell surface markers than healthy donors (supplemental Figure 11A). Following gene-correction, the CD56^bright^CD16^-^ population was reduced, although not to the same levels observed in healthy donors (supplemental Figure 11B). These results show that gene-corrected HSPCs rescue both the adaptive and innate immune defects.

## DISCUSSION

We describe a proof-of-concept study for a novel therapy for *RAG2* deficiencies. Through a broad and diverse set of preclinical studies performed *in vitro, in vivo*, in immunodeficient mice, using healthy donor and *RAG2*^*null*^ patient-derived HSPCs, we have demonstrated the application of CRISPR/Cas9-AAV6 platform in correcting a *RAG2*^null^ patient’s hematopoietic stem cells to restore the V(D)J activity in support of lymphopoiesis. For primary immunodeficiencies (PIDs) like *RAG2*-SCID, where a strict spatiotemporal level of gene expression and regulation is required for normal immune development and function^3,43^, a gene-correction approach has several safety and therapeutic benefits when compared to semi-random gene addition strategy: (1) it restricts the gene activity to the lymphoid lineage and the G_0_-G_1_ cell cycle phase [safety], (2) it circumvents the risk of genotoxicity from ubiquitous *RAG* activity [safety], (3) it allows for physiological gene expression [efficacy], and (4) it is easily adaptable to correct other forms of *RAG2* deficiencies, such as OS, AS and CID-G/AI for which disrupting and correcting the mutant alleles both alleviate disease burden and achieves cure [efficacy].

While lentiviral (LV) gene therapy is being developed for RAG1-SCID^44^ (a closely but genetically distinct from *RAG2*-SCID) and the safety and efficacy of delivering the RAG1 nuclease in HSPCs are being tested (NCT04797260), no alternative therapies, other than allo-HSCT are currently available for patients with *RAG2* deficiencies. We report the first feasibility study of gene-correction at the *RAG2* locus in human HSCs derived from healthy and *RAG2*^null^ patients to assess the safety and efficacy of this strategy. Allo-HCT for *RAG1/2-*deficient patients shows clinical improvement if a minimum of 20% donor chimerism is achieved^11^, an allele correction threshold that we have achieved and surpassed. We demonstrate that co*RAG2*-HSPCs can support durable hematopoiesis (18-20 weeks in immunodeficient mice), with only a 2-fold decrease in engraftment potential of corrected *RAG2*^null^ patient’ cells. This decrease could be explained by a heightened sensitivity to non-GMP grade virus, an observation confirmed by the CFU assay (data not shown), or a reduced regenerative potential of patients’ cells. In support of the latter, we show that the unmodified *RAG2*^null^ HSPCs have a 4.5-fold lower engraftment capacity than healthy-donor HSPCs controls. Alternatively, in the absence of *RAG2* expression, patient cells with lower DNA repair capacity are not selected against, resulting in an overall decrease in the fitness of the hematopoietic progenitors (HPCs), necessary for supporting HSCs engraftment after conditioning^42^.

We demonstrate *in vivo* rescue of B-cell development following gene-correction, but no detectable IgG heavy chain transcripts in the corrected and wild type sorted mature B cells, possibly due to the absence of a foreign antigen challenge to stimulate class switch recombination (CSR) in immunodeficient mice housed under pathogen-free conditions. When translating results from humanized mouse models to clinical settings, we acknowledge the microbiome differences between mice and humans and the critical role it plays in shaping the immune system’s development and function^45^. Humanized mice models have an underdeveloped immune system, which could underestimate the potency of the therapeutic product we are testing.

Gene-targeted *RAG2*^null^ HSPCs allowed correction of the peripheral T cell compartment, as documented in the spleen of transplanted mice. This was especially evident when a higher cell dose was transplanted into mice. It is possible that with lower dose cells, fewer corrected early thymic progenitors (ETP) survive the mouse thymic selection in support of lymphopoiesis. Delivering a higher dose of corrected cells could compensate for this limitation. We show that the ETP that survived the thymic selection developed into mature SP CD4^+^ and CD8^+^ T lymphocytes expressing αβ γδ T-cell receptors that can propagate developmental signals, supporting single-positive CD4^+^ and CD8^+^ T-cells. The newly derived receptors have an oligoclonal rearrangement pattern at the *TRA/TRD* locus with a similar pattern in usage of *TRAV* and *TRAJ* genes to those derived from healthy controls. This contrasts with the previous findings, where we demonstrated polyclonal TCR repertoire in T-cells, derived *in vitro*, from gene-corrected *RAG2*^null^ HSPCs^31^. It is reasonable to assume that of the gene-corrected and transplanted HSPCs clones, only a limited number repopulated the mouse thymus, thus restricting the number of stem cells that can contribute to the T-cell lineage development. This is supported by a recent study whereby using lentivirus cellular barcoding and purified human BM- or CB-derived HSPCs transplanted into NSG mice, less than 10 HSC clones contributed to the repopulation of the mouse thymus microenvironment^46^. Although the study further showed that even with a limited clonal contribution to the T-cell lineage, a diverse and polyclonal TCR repertoire can be generated^46^, we reason that the fresh CB- or BM-derived HSPCs used in their study holds greater lymphoid potential than frozen PB healthy or patient-derived HPSCs used in the study.

In the absence of myeloablative conditioning, *RAG*-deficient patients show a high graft failure rate, and those who engraft often have poor lymphoid reconstitution^11^. It has been speculated that genetically defective early CLPs occupy the bone marrow and thymic niches^16^. In the absence of niche clearing, the donor functional CLPs must compete with the endogenous genetically defective progenitors for space to engraft and develop. Moreover, we and others have recently shown that CD34^+^ cells from *RAG*^null^ patients can differentiate up to CD4^+^CD8^+^ double-positive (DP) cells in vitro^47,48^, suggesting that after HCT, competition between host and donor-derived cells might exist up to the DP stage. Additionally, it has been reported that the *RAG*-patient derived NK cells have higher perforin expression and increased degranulation potential^41,49^. These cytotoxic NK cells can attack the graft in the absence of conditioning. We show that there is a shift in the predominant NK population following gene-correction from a CD56^bright^ CD16^-^ to a CD56^dim^ CD16^+^. We thus propose that an autologous HCT using gene-corrected HPSCs, combined with reduced intensity myelosuppressive conditioning alone or in combination with antibody-based depletion, such as anti-CD45 ^50,51^, would conceivably provide therapeutic benefits by eliminating the competitive environment in the hematopoietic niches and assuring graft survival through correction of the cytotoxic function of the derived NK cells.

Finally, using i53 treatment resulted in ∼1.5-fold higher gene-targeting (HR-GT) without altering engraftment potential. We anticipate that a GMP-grade virus would decrease cellular toxicity and eliminate the use of i53, simplifying the path to clinical translation. Our studies provide the rationale for further testing our strategy to correct the immune dysregulations in *RAG2-*deficient patients.

## ACKNOWLEDGMENTS

We thank Stanford’s Binns Program for Cord Blood Research and patients for providing an invaluable source of CD34^+^ HSPCs for research. We thank the Stanford FACS Core facility for providing technical support. We acknowledge BioRender graphic software for helping with high-quality schematics. M.P-D gratefully acknowledges the funding support from the Primary Immune Deficiency Treatment Consortium (U54 AI 082973), part of the NCATS Rare Diseases Clinical Research Network (RDCRN), an initiative of the Office of Rare Diseases Research (ORDR), NCATS. The PIDTC is funded by a collaboration between NCATS and the National Institute of Allergy and Infectious Diseases (NIAID). M.H.P gratefully acknowledges the support from the Amon Carter Foundation and the Lacob Endowment. L.D.N gratefully acknowledges the funding support from the Division of Intramural Research (NIH/NIAID). We thank members of the Porteus Laboratory for valuable discussions and support.

## AUTHORSHIP

M.P.-D. and C.L.G, designed and performed most of the experiments. Y.G.N carried out the IF engraftments with *RAG2*-SCID patient cells and, together with R.M., provided thoughtful comments on the manuscript. T.K. carried out the BCR analysis. O.M.D., B.P., M.B., and F.P. carried out TCR analysis. S.V. assisted with mouse studies and provided thoughtful comments on the manuscript. H.G. assisted with AAV6 virus production and testing. N.B., G.L., and C.V. carried out the off-target analysis, designed and purified i53 protein. A.S. and T.M. obtained and purified CD34^+^ HSPCs from donated cord blood. J.B and E.M.H.G assisted with CFU assays. H.L.M obtained patient cells and provided thoughtful comments on the manuscript. L.D.N. and M.H.P. supervised the work, assisted with experimental design, and provided thoughtful feedback and discussion of the manuscript. M.P.-D., L.D.N., and M.H.P are responsible for all the data.

---

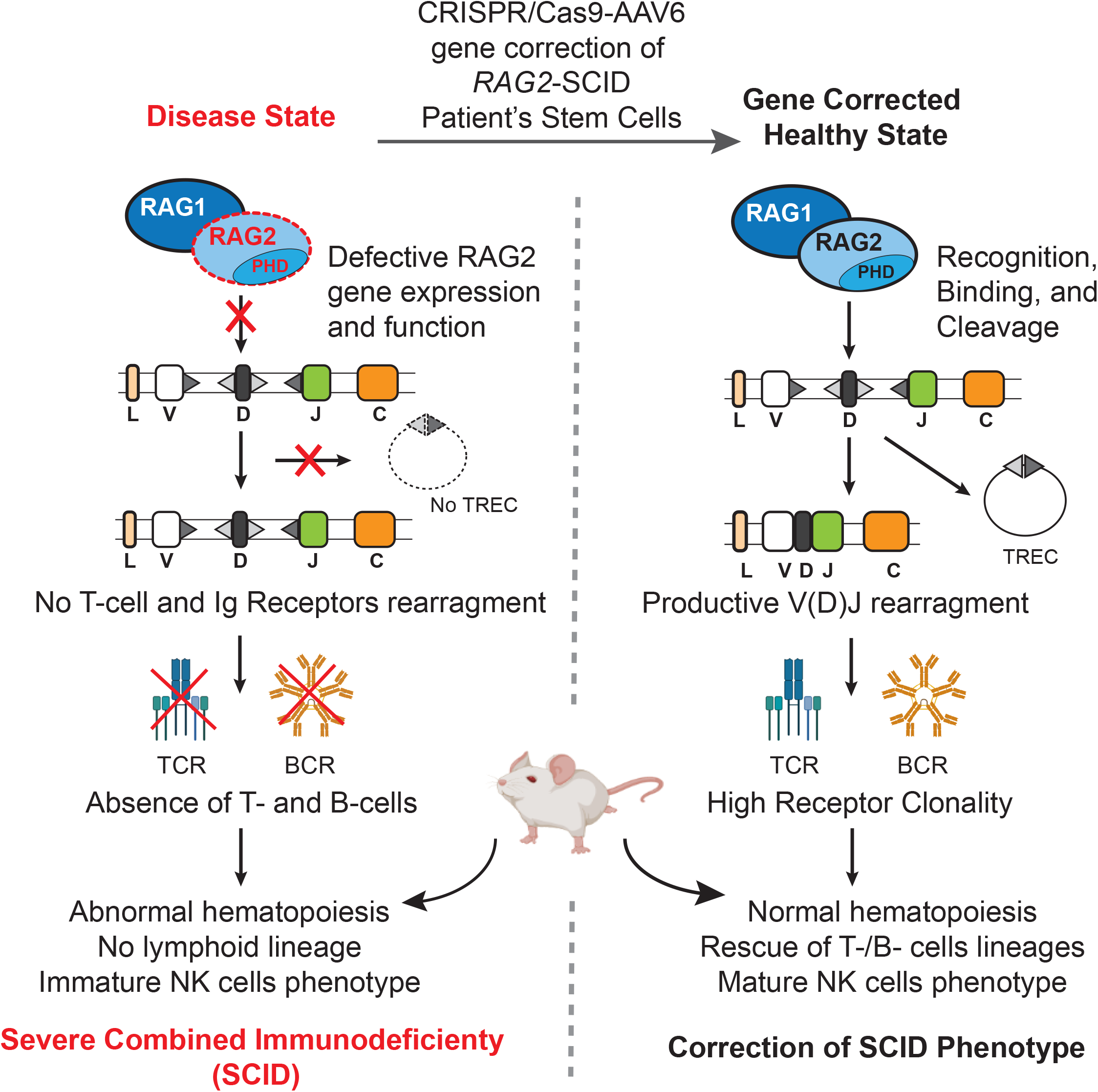

