## Supplemental Figure 1 for "Genetically Corrected *RAG2*-SCID Human Hematopoietic Stem Cells Restore V(D)J-Recombinase and Rescue Lymphoid Deficiency"

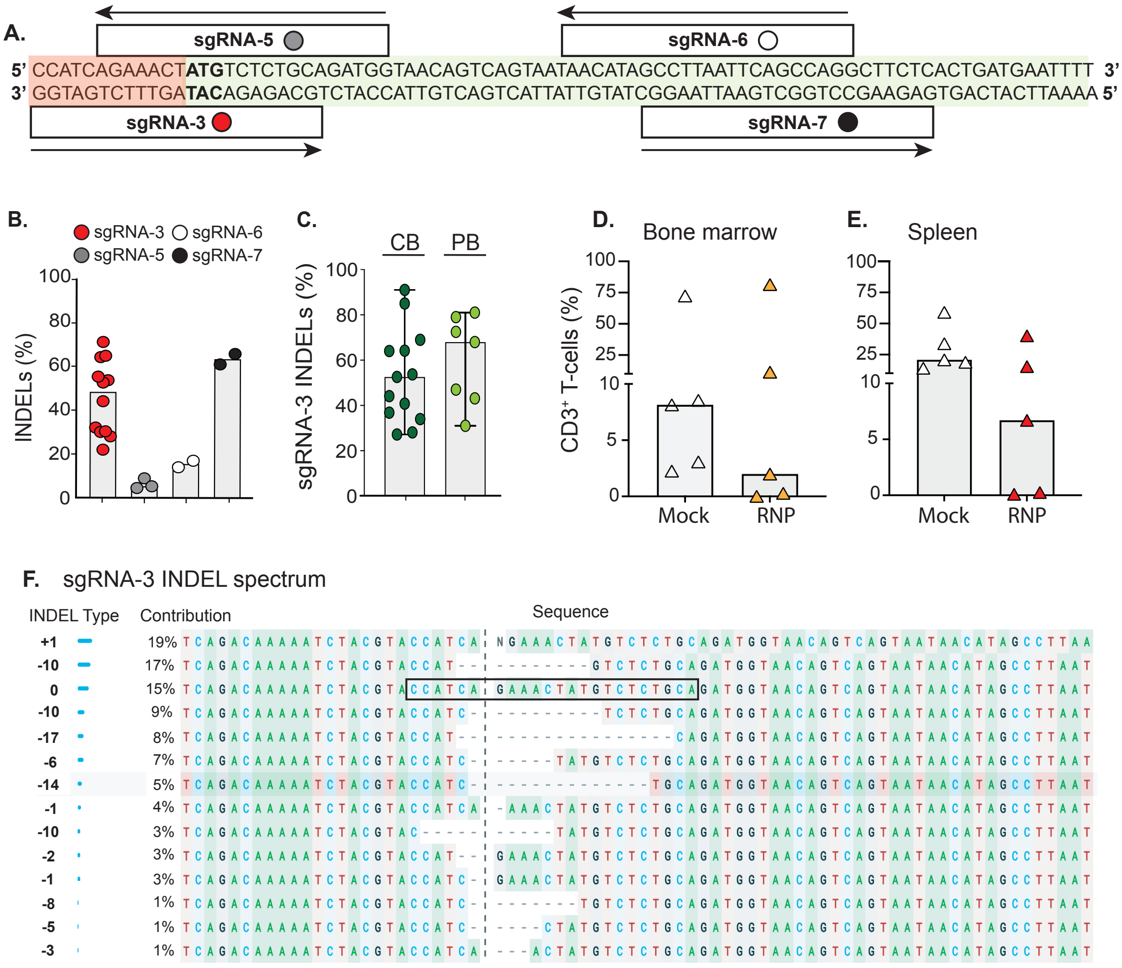


**Supplementary Figure 1. Characterization of RAG2 sgRNA-3. (A)** Schematic of sgRNAs (1-4) binding sites at the RAG2 genomic locus. Orange highlighted sequence indicates the untranslated region and the green highlighted sequence marks the RAG2 gene open reading frame, starting from the transcription start site (ATG, marked in bold). **(B)** Assessing genome editing efficiency (INDELs frequency) with the indicated RAG2 sgRNAs. INDELs (insertion and deletions) quantified 48h post-nucleofection by Sanger sequencing. **(C)** Percent INDELs generated by sgRNA-3 in cord blood (CB) and peripheral blood (PB)-derived HSPCs. **(D)** FACS-based quantification of CD3+ T-cells at week 18, post-Tx into immunodeficient mice, using healthy donor-derived UC HSPCs. A 4-fold decrease in bone marrow (BM) and **(E)** 3.1-fold decrease in the spleen (SP) in $CD3^{+}$ T cells derived from RNP treated conditions (orange, BM and red triangle, BM) as compared to mock-treated control (white triangles). Each triangle represents an individual mouse. **(F)** INDEL spectrum generated by RAG2 sgRNA-3. 82% of alleles acquired INDELS at 48 hours post-RNP-treatment using HiFi Cas9 nuclease. INDEL types and combinations are shown. sgRNA-3 sequence and PAM site are boxed. Analysis was done using Synthego ICE software. Bars: median.
