## Supplemental Figure 2 for "Genetically Corrected *RAG2*-SCID Human Hematopoietic Stem Cells Restore V(D)J-Recombinase and Rescue Lymphoid Deficiency"

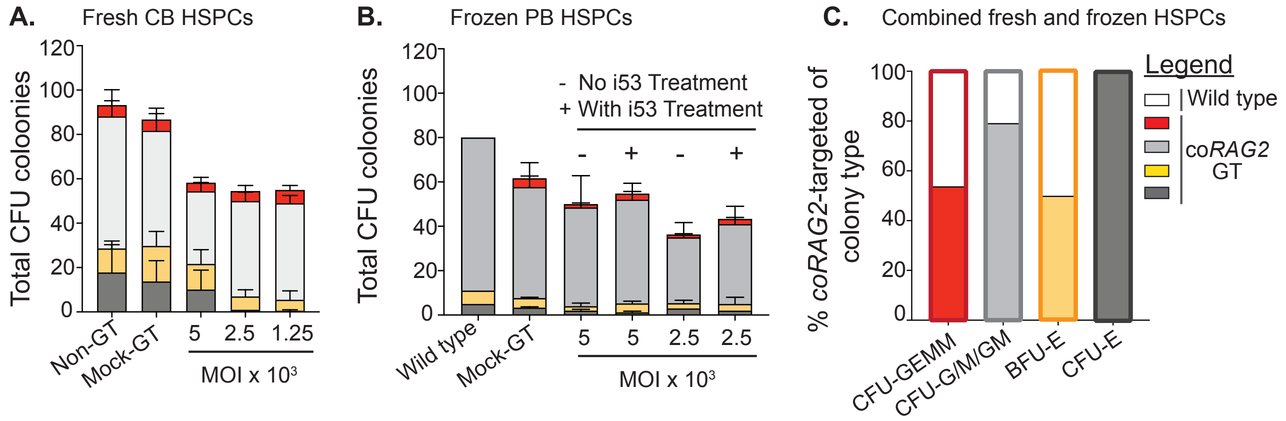


**Supplementary Figure 2. Colony-forming units (CFU) potential of coRAG2 gene-targeted HSPCs.** **(A)** Colony-forming units (CFU) derived from coRAG2-GT CB and **(B)** PB-derived HSPCs. Non-gene targeted (non-GT, wild type HPSCs), mock GT (nucleofected only), or GT with coRAG2 cDNA at various AAV6 multiplicity of infection (MOIs). Multi-potential granulocyte, erythroid, macrophage, megakaryocyte progenitor cells (CFU--GEMM, red) granulocytes, macrophage, or both (CFU-G/M/GM, light grey), erythroid progenitors burst-forming unit (BFU-E, yellow), and CFU erythroid (CFU-E, dark grey). n = 4 unique CB-derived HPSC donors for WT, mock, and GT using 5,000 MOI. n = 2 unique PB-derived HSPCs donors for GT using 2,500 MOI and 1,250 MOI. HSPCs treated (+) or not (-) with p53 inhibitor (i53). **(C)** Percent coRAG2-targeted alleles (colored bars) and non-targeted alleles (white bars) for each CFU type. Quantification by ddPCR. 360 total colonies scored.
