## Supplemental Figure 3 for "Genetically Corrected *RAG2*-SCID Human Hematopoietic Stem Cells Restore V(D)J-Recombinase and Rescue Lymphoid Deficiency"

**A. CFU analysis.** Biological replicate 1.

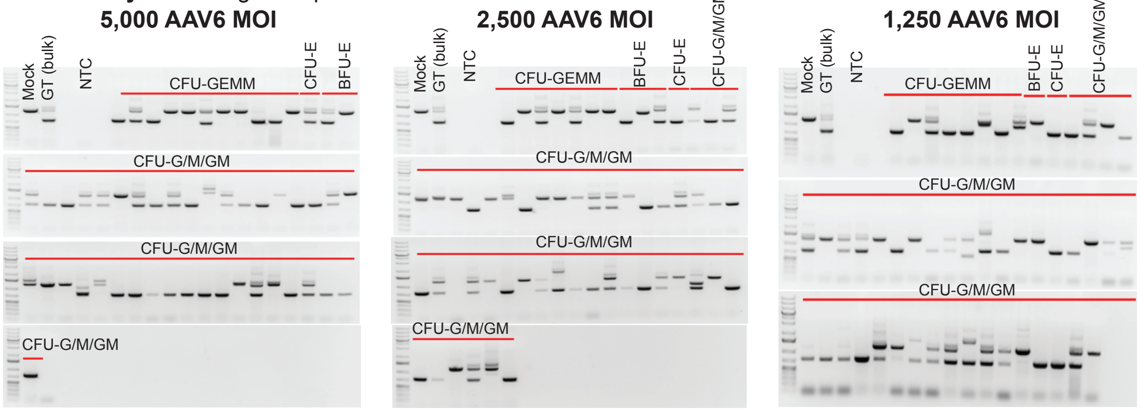


**Table 1. Percent of total CFU colony types derived from the indicated virus MOI.** Biological replicate 1.

| Colony Type | % *RAG2*-GT: 5,000 MOI | % *RAG2*-GT: 2,500 MOI | %*RAG2*-GT: 1,250 MOI |
|---|---|---|---|
| CFU-E | 100.0 | 66.7 | 100.0 |
| BFU-E | 0 | 100.0 | 0 |
| CFU-G/M/GM | 74.5 | 66.7 | 60.1 |
| CFU-GEMM | 54.5 | 42.8 | 62.5 |

**B. CFU analysis.** Biological replicate 2.

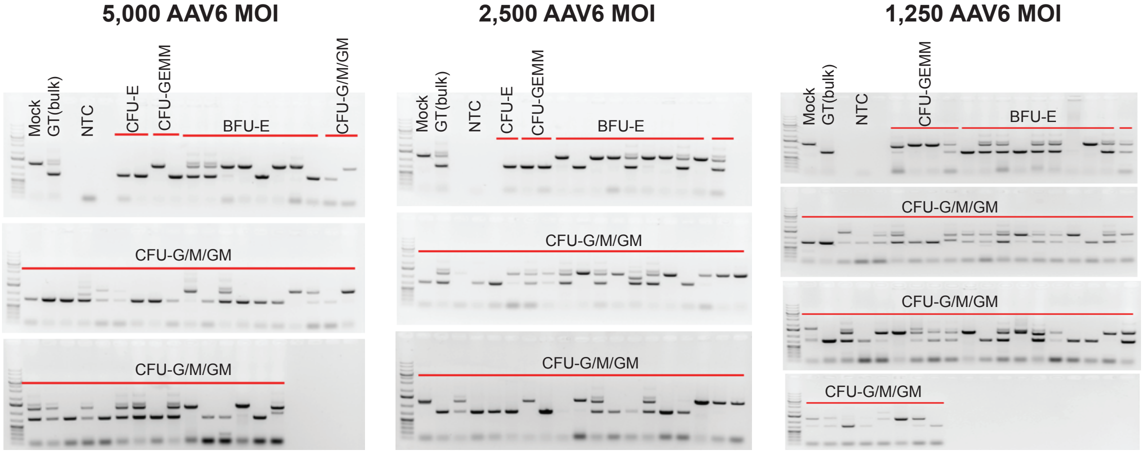


**Table 2. Percent of total CFU colony types derived from the indicated virus MOI.** Biological replicate 2.

| Colony Type | % *RAG2*-GT: 5,000 MOI | % *RAG2*-GT: 2,500 MOI | %*RAG2*-GT: 1,250 MOI |
|---|---|---|---|
| CFU-E | 100.0 | 100.0 | 0 |
| BFU-E | 50.0 | 100.0 | 77.8 |
| CFU-G/M/GM | 78.4 | 33.3 | 80.8 |
| CFU-GEMM | 50.0 | 61.5 | 25.0 |

**Supplementary Figure 3. Distribution of mono- and bi-allelic modification in coRAG2 gene-targeted healthy donor-derived HSPCs. (A)** coRAG2 gene-targeted CB-derives HSPCs from donor 1 and **(B)** donor 2 using three different AAV6 MOIs were single cells FACS sorted into one well of a 96-well plate coated with methylcellulose containing human stem cells specify cytokines. CFU colonies were genotyped 14-days post--sorting. A three primer "in-out" PCR reaction was developed to distinguish and quantify the alleles carrying the coRAG2 cDNA from the wild-type alleles.
