## Supplemental Figure 4 for "Genetically Corrected *RAG2*-SCID Human Hematopoietic Stem Cells Restore V(D)J-Recombinase and Rescue Lymphoid Deficiency"

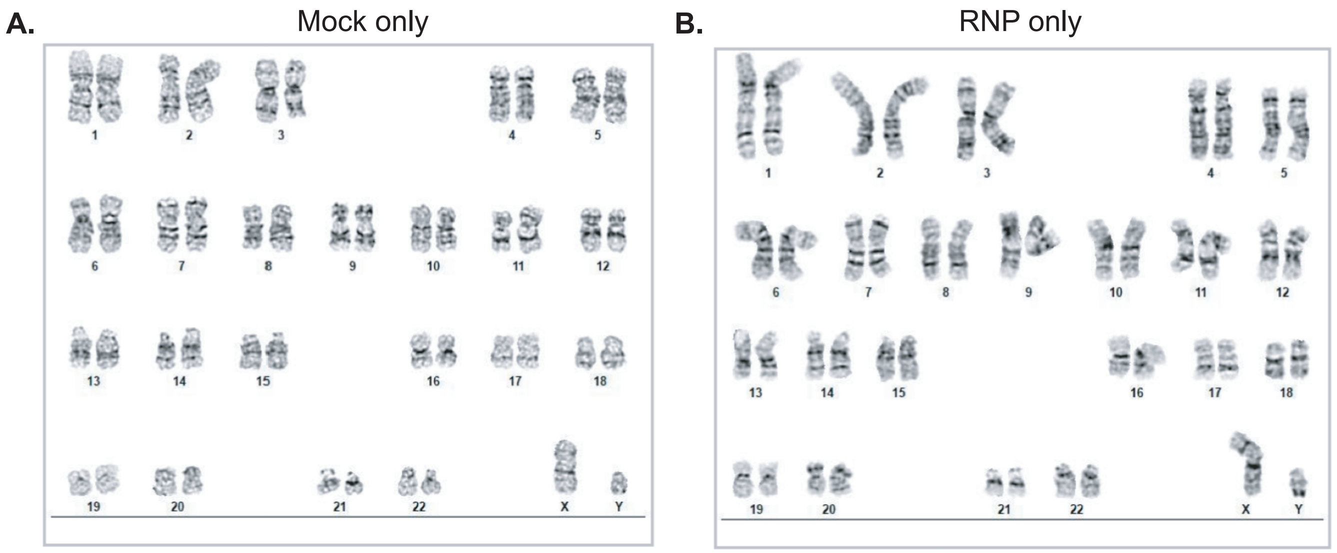


**Supplementary Figure 4. Karyotype analysis post-RNP treatment in RAG2null patient-derived HSPCs.** **(A)** RAG2-SCID (c.296C>A; c.1342C>A) HSPCs were nucleofected alone or **(B)** with sgRNA-3 and HiFi Cas9 nuclease (RNP). G-band karyotype analysis was carried out on 20 cells per condition. No clonal abnormalities were detected at the 375-425 band resolution for the mock-treated and at 350-400 for the RNA-treated samples.
