## Supplemental Figure 5 for "Genetically Corrected *RAG2*-SCID Human Hematopoietic Stem Cells Restore V(D)J-Recombinase and Rescue Lymphoid Deficiency"

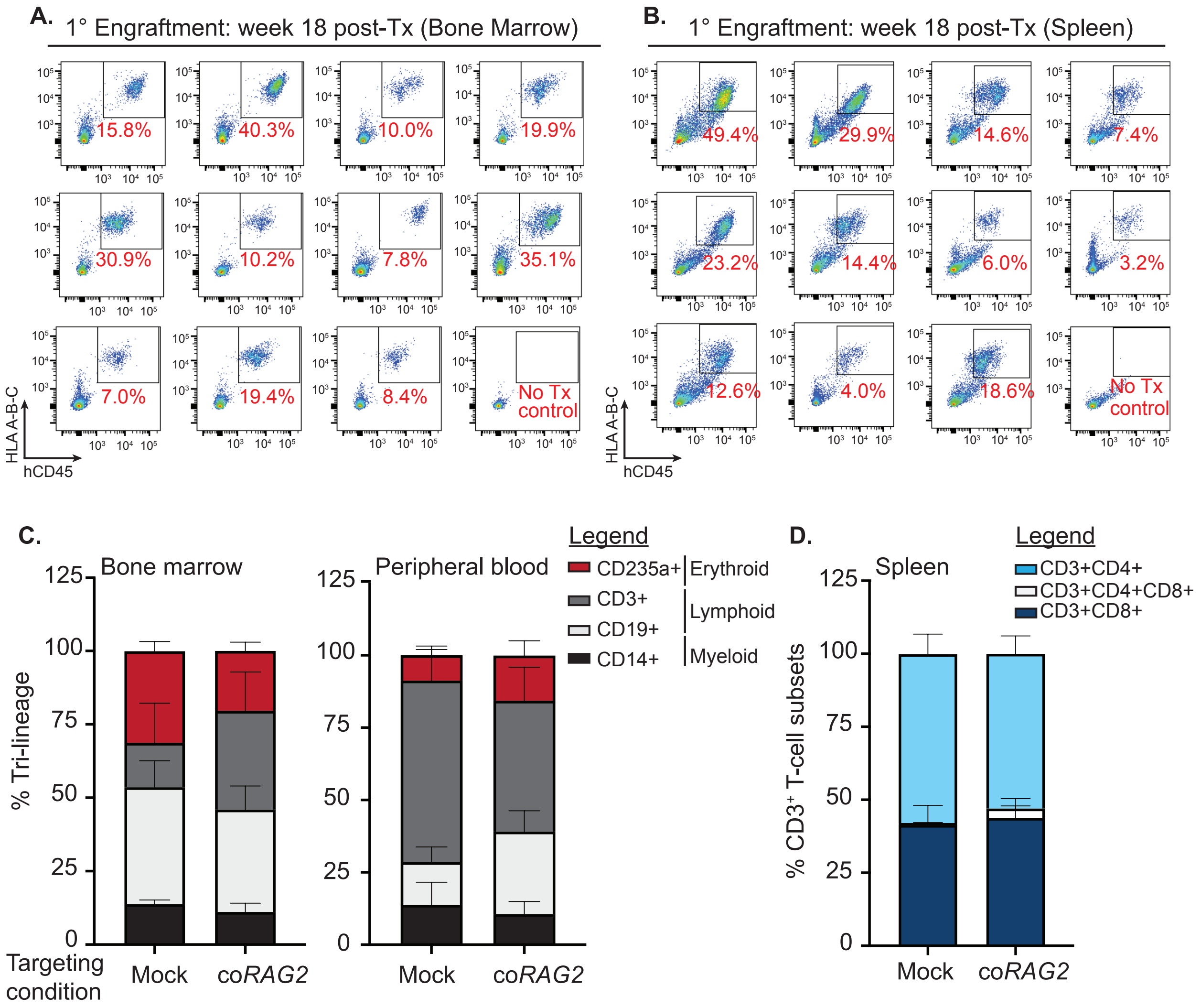


**Supplementary Figure 5. In vivo long-term human hematopoietic reconstitution from coRAG2-targeted healthy donor-derived HSPCs. (A)** FACS-based quantification of bone marrow (BM) and **(B)** spleen (SP) primary human engraftment (hCD45$^{+}$ HLA A-B-C$^{+}$) derived from coRAG2-targeted fresh CB HSPCs. Each FACS plot represents an individual mouse. **(C)** Endpoint analysis (22 weeks post-Tx) of human hematopoietic lineage distribution in the BM and peripheral blood of immunocompetent mice (NSG). CD235$^{+}$ (erythroid), CD3$^{+}$ (T-cells), CD19$^{+}$ (B-cells), CD14$^{+}$ (monocytes); n=5 mice (mock), n=11 (coRAG2-GT). **(D)** CD3$^{+}$ T cell subsets (CD4$^{+}$, CD8$^{+}$, CD4$^{+}$CD8$^{+}$) derived in the spleen of NSG mice from (B) engrafted with coRAG2-GT HSPCs.
