## Supplemental Figure 6 for "Genetically Corrected *RAG2*-SCID Human Hematopoietic Stem Cells Restore V(D)J-Recombinase and Rescue Lymphoid Deficiency"

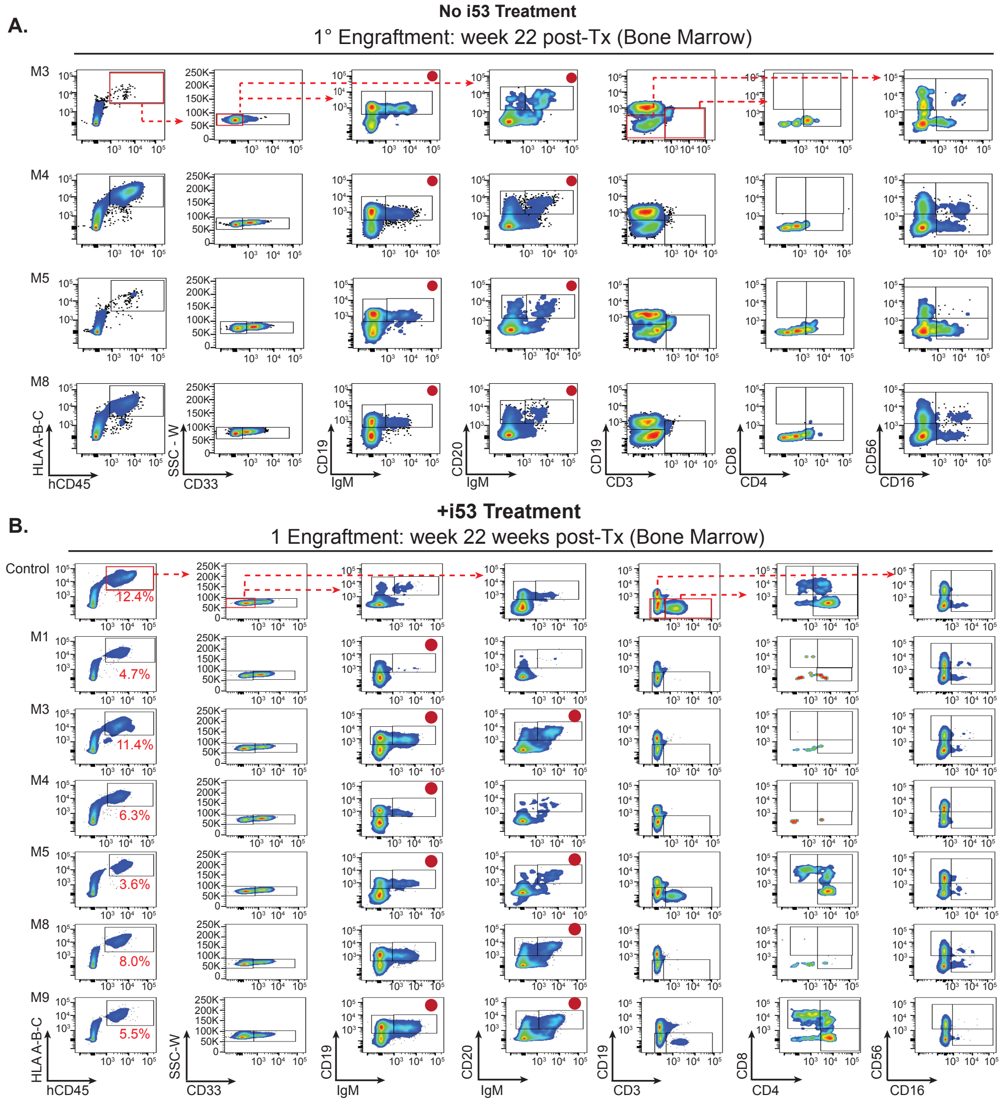


**Supplementary Figure 6. In vivo human hematopoietic reconstitution derived from coRAG2-GT healthy donor HSPCs. (A)** Lymphoid lineage reconstitution derived from coRAG2-GT HSPCs not treated with i53 inhibitor. The gating strategy is shown. Marked FACS plots (red dot) indicate the cell type sorted in Figure 3E. **(B)** same as (A) but treated with i53 (+i53) inhibitor. Control: wild-type HSPCS engrafted into immunodeficient NSG mice. M: mouse ID; $CD45^+$ HLA A-B-$C^+$: double-positive human engraftment quantified by FACS in the mouse BM; CD33: separates myeloid ($CD33^+$) from non-myeloid ($CD33^-$) human hematopoietic lineages; $CD19^+$ and $CD20^+$: B-lymphocytes; CD3+: pre-T lymphocytes; $CD3^+CD4^+$: T-helper cells; $CD3^+CD8^+$: T-effector cells; $CD3^-CD16^+$, $CD3^-CD56^+$ and $CD3^-CD16^+CD56^+$: subsets of natural killer (NK) cells.
