## Supplemental Figure 7 for "Genetically Corrected *RAG2*-SCID Human Hematopoietic Stem Cells Restore V(D)J-Recombinase and Rescue Lymphoid Deficiency"

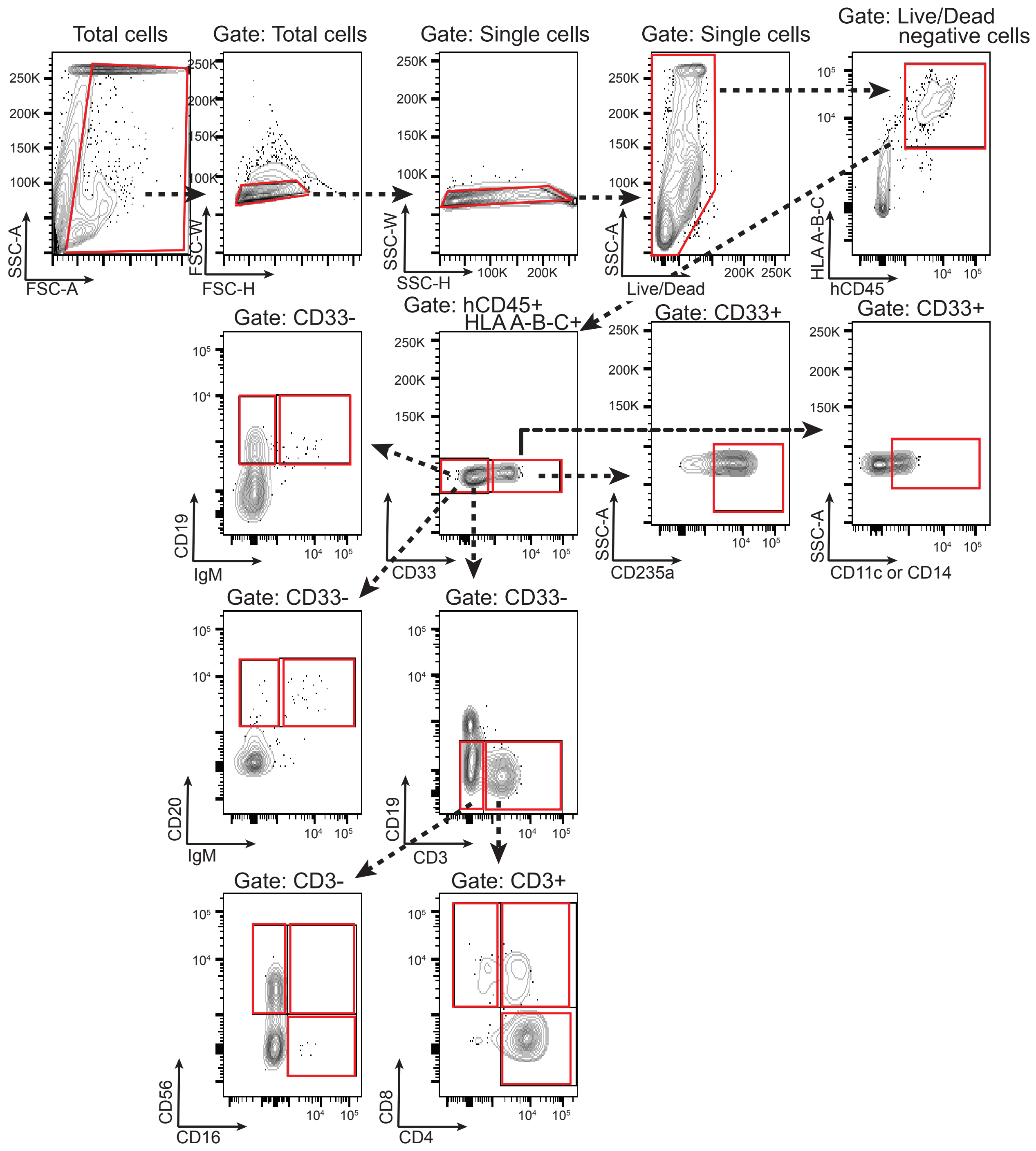


**Supplementary Figure 7. Human engraftment and tri-lineage gating scheme.** Gating strategy was used for data presented in Figures 3A, 3B, 3E, 3F, 4B, 4E, 5A-C, S5, S6.
