## Supplemental Figure 8 for "Genetically Corrected *RAG2*-SCID Human Hematopoietic Stem Cells Restore V(D)J-Recombinase and Rescue Lymphoid Deficiency"

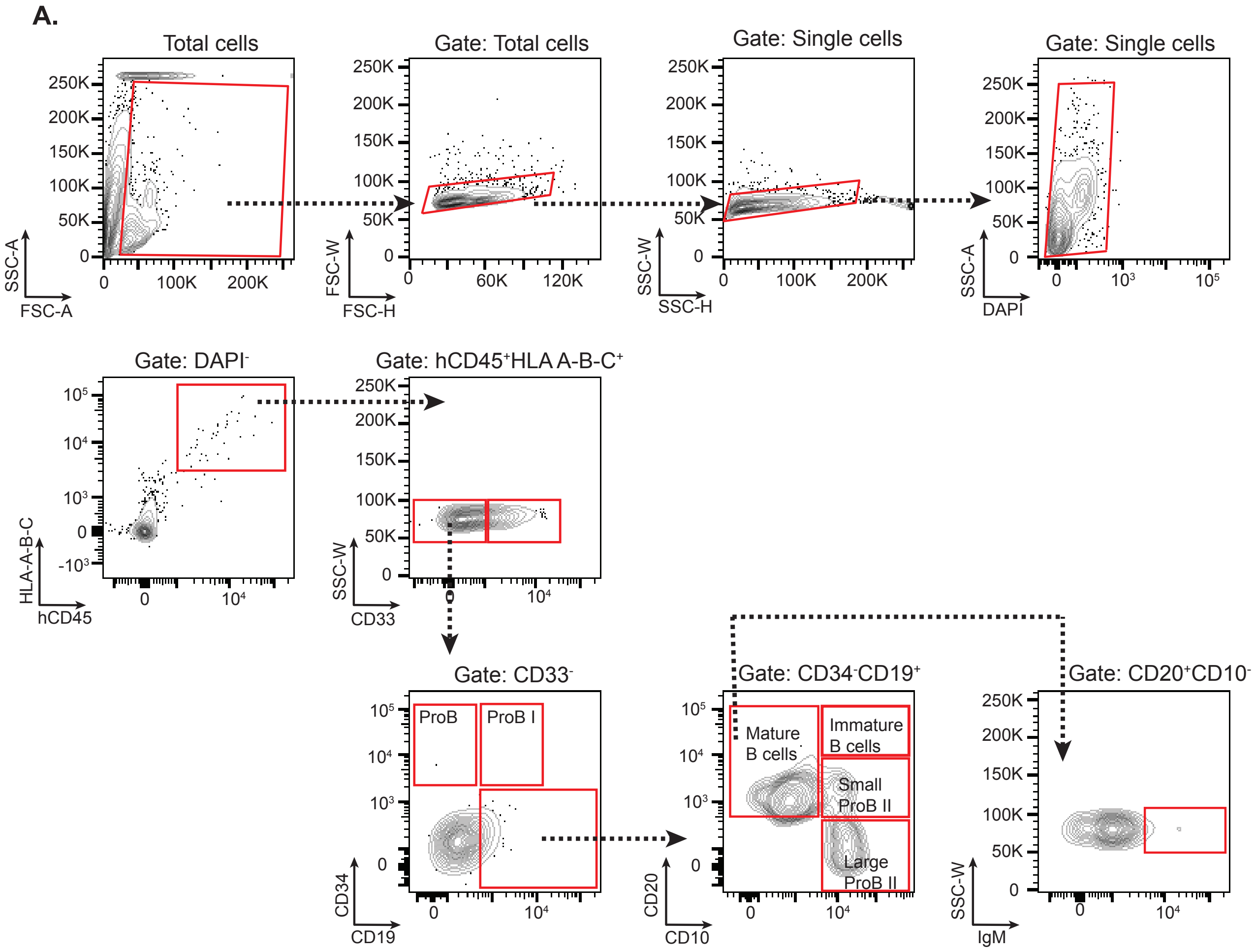


**Supplementary Figure 8. B-lymphoid development from coRAG2-GT of RAG-SCID patient-derived HSPCs.** The gating strategy is used for data presented in Figures 5A-5E.
