## Supplemental Figure 9 for "Genetically Corrected *RAG2*-SCID Human Hematopoietic Stem Cells Restore V(D)J-Recombinase and Rescue Lymphoid Deficiency"

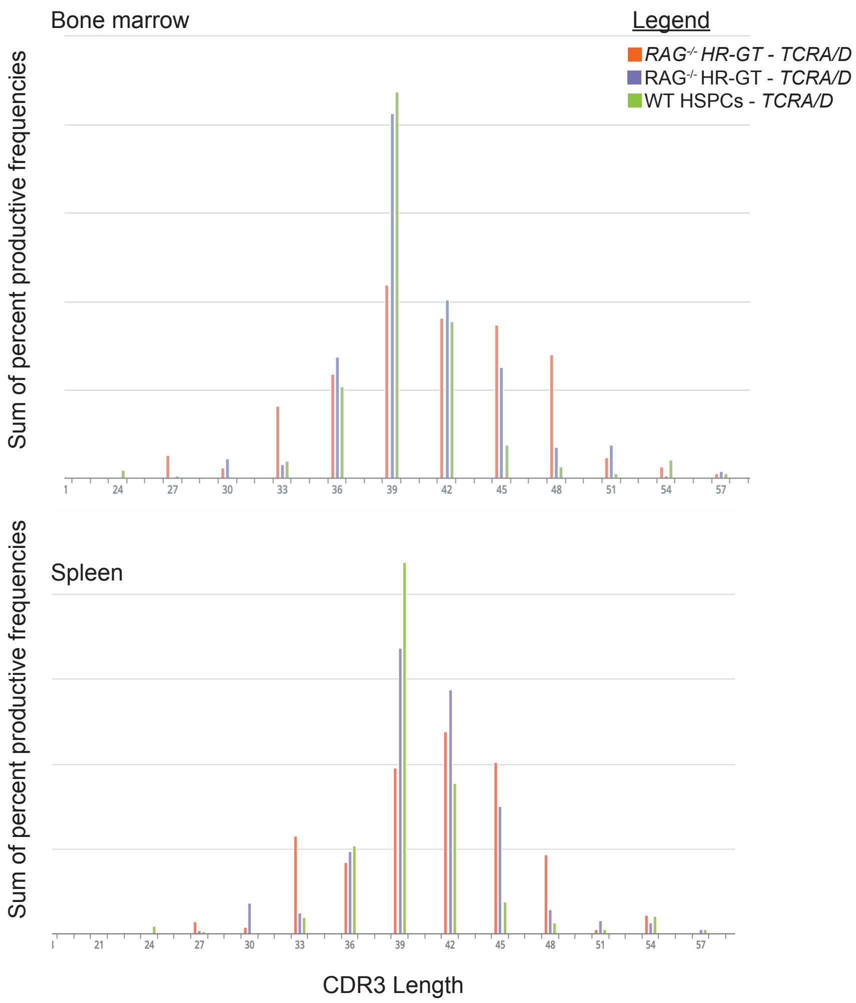


**Supplementary Figure 9. Virtual spectratyping of RAG2null patient and healthy donor- HSPCs derived mature T-cells.** Similar CDR3 lengths in TCRAD chains were obtained from the pools of wild type (healthy donor) and genome-targeted patient T cells expressing $CD3^{+}$.
