## Supplemental Figure 10 for "Genetically Corrected *RAG2*-SCID Human Hematopoietic Stem Cells Restore V(D)J-Recombinase and Rescue Lymphoid Deficiency"

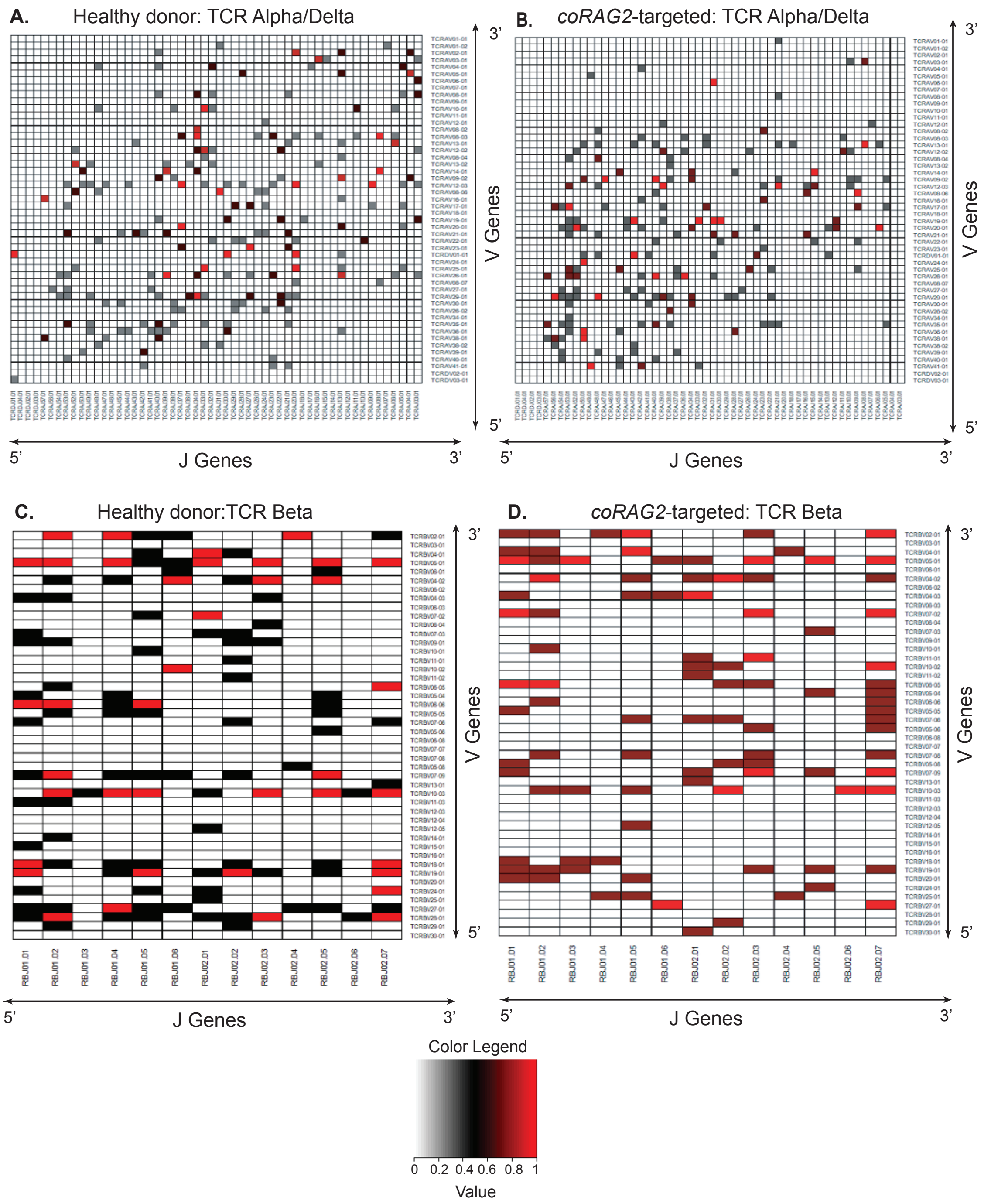


**Supplementary Figure 10. Heatmaps of V and J genes pairing for TCR A/D and TCR beta of T-lymphocytes sorted from gene-targeted (GT) RAG2null patient-derived HSPCs and engrafted into immunodeficient mice. (A)** Heatmaps of V and J genes pairing for TCR A/D in healthy donors. **(B)** coRAG2-GT or for **(C)** TCR B in healthy donors and **(D)** coRAG2-GT and derived $CD3^{+}$ sorted cells. V and J genes are listed in the order of their chromosomal location.
