## Supplemental Figure 11 for "Genetically Corrected *RAG2*-SCID Human Hematopoietic Stem Cells Restore V(D)J-Recombinase and Rescue Lymphoid Deficiency"

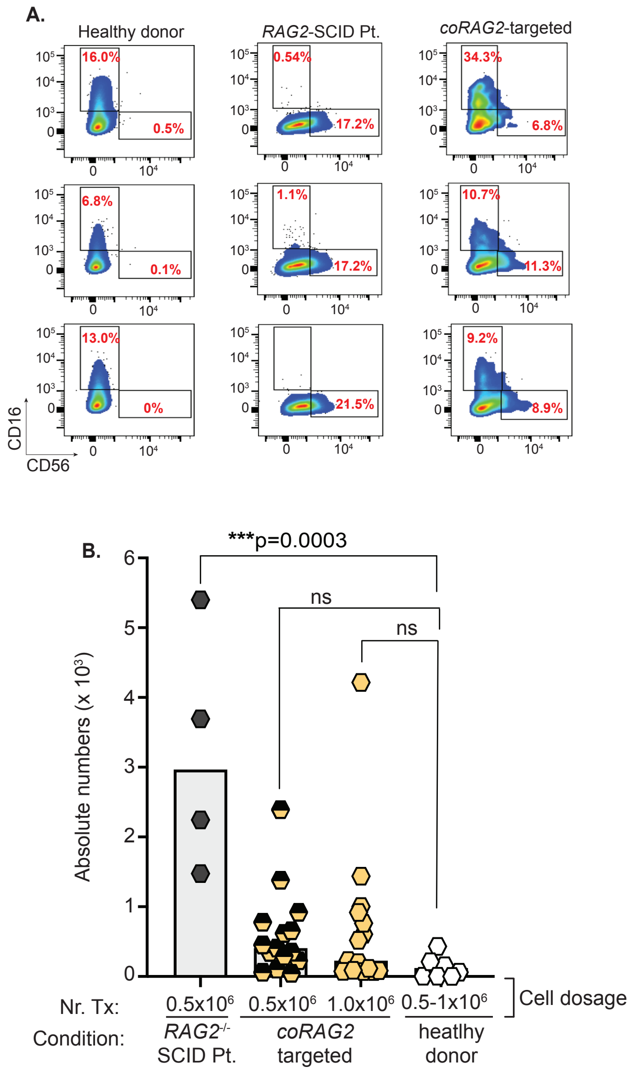


**Supplementary Figure 11. Phenotypic correction of NK-lymphocytes defect in RAG2null patient--derived HSPCs.** (A) FACS plots showing human NK-cells development in the bone marrow of immunodeficient mice (22 weeks post-Tx). Cells gated on CD3-. (B) The absolute number of NK cells derived from RAG2$^{null}$ patients (n=4 mice, black diamond), coRAG2-GT patient cells (n=15, half diamonds; n=15, yellow diamonds), and healthy donors (control, n=7, white diamonds). Stats: one-way ANOVA, nonparametric, Dunn’s multiple comparison tests. Bars: mean ± s.e.m.
