## Supplemental Material and Methods for "Genetically Corrected *RAG2*-SCID Human Hematopoietic Stem Cells Restore V(D)J-Recombinase and Rescue Lymphoid Deficiency"

Short Title: **Gene correction approach for *RAG2*-SCID deficiency**

Mara Pavel-Dinu[1†], Cameron L. Gardner[2,3†], Yusuke Nakauchi[4], Tomoki Kawai[2], Ottavia M. Delmonte[2], Boaz Palterer[2], Marita Bosticardo[2], Francesca Pala[2], Sebastien Viel[1,5], Harry L. Malech[6], Hana Y. Ghanim[1], Nicole M. Bode[7], Gavin L. Kurgan[7], Christopher A. Vakulskas[7], Adam Sheikali[1], Sherah T. Menezes[1], Jade Chrobok[1], Elaine M. Hernández González[1], Ravindra Majeti[4], Luigi D. Notarangelo[2*], Matthew H. Porteus[1*]

### Supplemental Methods

**Gene-editing (INDELs) and gene-targeting (HR) quantification.** Insertions and deletions (INDELs) frequencies were quantified using TIDE or ICE (Synthego) online software on genomic DNA extracted using Quick Extract, Epicentre (Madison, WI, USA), and amplified using F (5' ATGTGGTTCTTTCAGCTG 3') and R (5' CGAAAAGTAACCTTTTTGTTGT 3') primers. Genomic integration was quantified by droplet digital PCR (ddPCR), as we previously described[26].

**sgRNA guide and HiFi *s.p.* Cas9.** Candidate *RAG2* sgRNAs were selected using CRISPOR software (crispor.tefor.net) and synthesized by Synthego Corp (Redwood City, CA, USA) as previously described[26]. Lead *RAG2* guide (RAG2-sg3: 5'-TGCAGAGACATAGTTTCTGA-3') cuts six base pairs upstream of the ATG translation initiation site. It was purchased from TriLink BioTechnologies (San Diego, CA, USA), as 2' O-methyl 3'phosphorothioate and HLPC-purified. HiFi *s.p.*Cas9 protein was purchased from IDT (Coralville, IA, USA).

**Off-target activity assay.** PCR amplicons were sequenced on an Illumina MiSeq instrument (v2 chemistry, 150bp paired-end reads) (Illumina). Data were analyzed and editing quantified using CRISPAltRations[1], utilizing the default parameters.

**Gene targeting (GT) based correction system.** The system uses a 2′-O-methyl 3′phosphorothioate chemically modified guide RNA (sgRNA) to direct a high fidelity (HiFi) Cas9 nuclease to a pre-defined genomic site where it creates double-strand breaks and activates the endogenous NHEJ and homologous direct repair (HDR) pathways. In the presence of adeno associated virus 6 (AAV6) that delivers a corrective donor DNA template carrying the desired genomic modification and flanked by homology arms to the break sites, the HDR pathway integrates the new DNA sequence through a homologous recombination-mediated genome targeting (HR-GT) process.

**Human stem cells isolation.** Frozen mobilized peripheral blood (PB) CD34$^+$ HSPCs were purchased from AllCells (Alameda, CA, USA) and thawed as previously described[1]. Fresh CB-derived CD34$^+$ HSPCs were obtained under informed consent from Binns Program for Cord Blood Research at Stanford University, purified, and cultured as previously described[26].

**Electroporation and transduction of human stem cells.** RNP complex was generated by mixing HiFi Cas9 with sgRNA at a molar ratio of 1:5 (450ug/ml HiFi Cas9 protein with 960 ug/ml of sgRNA from Trilink), at 37°C for 30 minutes prior to nucleofection. $1x10^6$ to $3x10^6$ HSPCs were nucleofected[26] in one well of a 16-well strip. Following nucleofection, cells were transduced, with AAV6 at 5,000 MOI unless otherwise stated. Mock control did not receive RNP complex.

**rAAV6 donor design and AAV6 virus purification.** The *RAG2* donor vector was constructed by PCR amplifying 400 bp left and right homology arms, flanking the RNP cut site, for the *RAG2* locus from human $CD34^+$ genomic DNA. BGH polyA and WPRE sequences were amplified from plasmids. Corrective, codon-optimized *RAG2* (co*RAG2*) cDNA was designed by GeneScript (Piscataway, NJ, USA) with silent mutations generating 76.3% homology to the endogenous gene. The codon-optimization was intended for reducing homology to the genomic locus and eliminating unwanted homologous recombination-based integration events while preserving the intact RAG2 amino acid sequences. While the co*RAG2* open reading frame disrupts the sgRNA binding site, the unmodified base pairs present in the 5'untranslated region, including the protospacer adjacent motif (PAM) sequence, remain intact. To assure that the integrated donor will not be re-cleaved, we introduced two mutations in the PAM sequence (CCA>GGA) located in the 5' homology arms of the donor. WPRE was included in the design to increase the transgene expression level if the codon optimization would reduce it. WPRE mechanism of action is not well understood, but it has been proposed to improve transcriptional termination or increase the efficiency of nuclear exports of transcripts. The donor plasmid was constructed using Gibson cloning New England Biolabs (Cat # E5510S) into a pAAV-MCS plasmid containing AAV2-specific inverted repeats (ITR) Stratagene (Santa Clara, CA, USA). AAV6 virus was purified, as we previously described[26].

**Expression and purification of i53 peptides.** A G-block gene fragment (Integrated DNA Technologies, Inc) encoding *E. coli* codon-optimized i53 peptide[2] containing an amino-terminal hexahistidine tag was subcloned into pET28b (Novagen). *E. coli* Bl21(DE3) expressing i53 was grown to mid-log phase at 37 °C ($OD_{600}$–0.6), at which time the culture was chilled to 30 °C and 1 mM isopropyl β-D-1-thiogalactopyranoside (IPTG) was added. Following shaking incubation (250 r.p.m.) at 30 °C for 3 h, cells were collected by centrifugation and lysed at 4 °C with the Emulsiflex-C3 high pressure homogenizer (Avestin, Ottawa ON, Canada). i53 peptide was purified from clarified lysates using immobilized metal affinity chromatography (IMAC), heparin chromatography, and size-exclusion chromatography protocols that have been described for i53 previously[2]. Purified proteins in this study were rigorously studied through commercial-grade analytical and quality control procedures. Briefly, proteins were analyzed for purity via denaturing SDS-PAGE and were determined be > 95% pure. Every protein was quality controlled to contain < 10 EU/mg of endotoxin, to be free of residual host cell DNA beyond the limits of detection with qPCR, and to be free of contaminating DNase and RNases.

**Absolute quantification of genome targeting (HR-GT).** To detect insertions at the *RAG2* genomic locus we used F- 5' TCT CAC CTC CCA TTC CCT AG 3', R - 5' TCA GGG CGA TAT TGT TGG AC, and labeled probe F – 5' FAM/CCC GTC TAG/ZEN/ TCA CTT CGC ACC TTC GGC/3IABkFQ 3'. The reference assay designed to detect the RAG1 reference genome sequence is: F-5' GCACAGGAAGTTTAGCAGTG 3', R-5'GGGAATTCAAGACGCTCAGA 3', and probe 5' HEX/CCC GAG GAA/ZEN/CGT GAC CAT GGA GTG GC/3IABkFQ 3'. Final concentration of primer and probes was 900 nM and 250 nM, respectively. The following ddPCR program was optimized to amplify a 516 bp and 512 bp for the targeted and reference amplicon, respectively: 1 – 95ºC for 10 min, ramp 1 ºC/sec, 2 – 94ºC for 30 sec, ramp 1 ºC/sec, 3 – 61.7ºC for 30 sec, ramp 1 ºC/sec, 4 – 72ºC for 2 min, ramp 1 ºC/sec, 5 – repeat steps 2-4 for 50 cycles, 6 – 98ºC for 10 min, ramp 1 ºC/sec, 7- 4ºC , ramp 1 ºC/sec. Bio-Rad Droplet Reader and QuantaSoft software were used generate and analyzed data, per manufacturer's guidelines. Absolute quantification (DNA/ul) was determined for reference and targeted genes. Total percent targeting was calculated as a ratio of HEX to FAM signal.

**Three-primer PCR-based genotyping of the colony-forming units (CFU).** PCR-based allele-specific genotyping of single-cell sorted-derived colonies (n = 900), as a function of virus multiplicity of infection (MOI). Primers used: F-WT: 5' TCACCTGTTCAAAAGTCCCC 3', R-integrated: 5' TGGTTGTGCTTCACGTCC 3', and R-WT 5'AGATGGTGTCATTTTTGGCAATAGAG 3'. The PCR reaction contained 0.5 uM of primers, 150-200 ng genomic DNA, and 1x Phusion Master Mix High Fidelity, per manufacturer's guidelines. The following PCR settings amplified the integrated band of 758 bp, a wild type band of 1246 bp: 1- 98ºC:30sec; 2-98ºC:10sec; 3-63ºC:30sec; 4-72ºC:30sec; 5-repeat septs 2-4 for 30 cycles, 6-72ºC:7min, 7-4ºC.

**Immunodeficient mice strains.** Two strains were used in the study: NSG (NOD.Cg-Prkdc$^{scid}$ IL2rg$^{tmlWjl/Sz}$) and NSG-SGM3 [NSG expressing stem cell factor (SCF), granulocyte-macrophage colony-stimulating factor (GM-CSF), and interleukin-3 (IL3). Animal work was carried out in pathogen-free conditions, in accordance with the Institutional Animal Care and Use Committee (IACUC) Approval, protocol number 25065.

**Patient *RAG2*-SCID HSPCs.** The patient was born at eight months of gestation and was well until 3 months of age when he developed rotavirus diarrhea, a diffuse skin rash, and eosinophilia. The skin rash was shown to be a manifestation of graft versus host disease due to maternal T cell engraftment. Genetic analysis performed on autologous cells revealed compound heterozygosity for two *RAG2* null variants (c.296C>A, c.1324C>A; p.P99Q, A442T). In vitro testing[1] of the recombination activities of these *RAG2* variants yielded values of 0% and 0.7% of wild-type *RAG2*, respectively, consistent with a diagnosis of *RAG2*-SCID.

**Transplantation of human genome modified CD34$^+$ HSPCs and engraftment assessment.** Human engraftment studies using fresh CB-HSPCs (2.5 x $10^5$) or frozen PB-HSPCs (5.0 x $10^5$) were injected intra-hepatic (IH) or intra-femoral (IF) into sub-lethally irradiated 2 days old NSG pups or 6-8 weeks old mice, respectively. Human chimerism defined as double-positive cells for hCD45$^+$ HLA A-B-C$^+$ out of total mouse cells was quantified 18-20 weeks post-engraftment in the bone marrow (BM) and spleen (SP) of mice injected with unmodified (wild type and mock), RNP-treated, and HR-GT HSPCs. The transplantation workflow was carried out as previously described[26] and in accordance with the approved Stanford University Administrative Panel on Lab Animal Care (APLAC).

**FACS analysis of human engrafted HPSCs.** All fluorescence activating cell sorting (FACS) analyses for human engraftment studies were done on FACS Aria II Sort Instrument part of the FACS Facility Core at Stanford University, Institute of Stem Cell Biology and Regenerative Medicine. The following antibody panel was used to analyze IF bone marrow engraftments. CD3-PerCP-Cy5.5 (clone: HiT3A, BioLegend); CD19-FITC (clone: HIB19, BioLegend); mCD45.1-PE Cy7 (clone:A20, BioLegend); CD10-PE Texas Red (clone:Hi10a, BD Biosciences); HLA A-B-C-APC-Cy7 (clone: W6/32, BioLegend); CD33-AF-700 (clone:WM53, eBioSciences); CD34-APC (clone:8G12, BD Pharmigen); hCD45-BV786 (clone: HI3A, BDHorizon); IgM-BV711 (clone:MHM-88, BioLegend); CD20-BV510 (clone:2H7, BioLegend); CD56-PacBlue (clone:MEM-188, BioLegend); CD16-BUV395 (clone:3G8, BD

Pharmigen); Live/dead (Invitrogen). The following antibody panel was used to analyze IF spleen engraftments: TCR a/b-PerCp-Cy5.5 (clone:iP26, BioLegend); TCR g/d-FITC (clone:cl.B1, BioLegend); CD45RA-PE Texas Red (clone: HI100, ); CD8-APC (clone:HiT8a, BioLegend); CD4 (clone:OKT4, BioLegend); CD3-BV421 (clone:UCHT1, BD BioSciences), mCD45.1, HLA A-B-C, CD33 and hCD45 same as above.

**Variable immunoglobulin M (IgM) heavy chain (Vh) B-cells analysis.** Total RNA was isolated using RNeasy Mini Kit (QIAGEN, Hilden, Germany) from $CD19^+$ B-cells sorted from the bone marrow of mice engrafted with human gene-targeted HSPCs. Total RNA was reverse transcribed with random primers using Superscript IV (Invitrogen, Carlsbad, CA, USA). cDNA was amplified using a 40 cycles PCR with primers specific for the seven variable heavy chains and with fluorescent reverse primer for the IgM isotype, as previously described[3]. PCR detection of variable immunoglobulin M (IgM) heavy chain (Vh). IgVH family: VH1 5′-TGGAGCTGAGSAGSCTGAGATCYGA-3′; VH2 5′-AACCCACASAGACCCTCAC-3′; VH3 5′-TCCCTKARACTCTCCTGTRCAGC-3′; VH4 5′-CTACAACCCSTCCCTCAAGAGT-3′; VH5 5′-CAGCACCGCCTACCTGCAGTGGAGC-3′; VH6 5′-TCCGGGGACAGTGTCTCT-3′; VH7 5′-CAGCACRGCATAYCTGCAGATCAG-3′; (IgH γ chain 5′-6Fam-AAGTAGTCCTTGACCAGGCAGC-3)′; IgH μ chain 5′-6Fam-GGAGACGAGGGGGAAAAGG-3′.

**High throughput sequencing (HTS) of TCRa and TCRb.** Bone marrow and spleen samples were collected at end-point engraftment analysis, as we previously described[4]. Genomic DNA was extracted using QIAamp DNA Micro Kit, Qiagen (Germantown, MD, USA). TCRa (*TRA*) and TCRb (*TRB*) rearranged genomic products were amplified by multiplex PCR (Adaptive Biotechnologies Seattle, WA) and analyzed using Adaptive Biotechnologies' assay-based computational techniques to minimize PCR amplification bias. The frequency of a given *TRA* or *TRB* sequence is representative of the frequency of that clonotype in the original sample. The PCR products were sequenced using the Illumina HiSeq platform. Custom algorithms were used to filter the raw sequences for errors and align the sequences to reference genome sequences. Subsequently, the data were analyzed using ImmunoSeq's online tools. The frequency of productive and nonproductive *TRA* or *TRB* rearrangements were analyzed within both unique and total *TRA* and *TRB* sequences obtained from T cells. The distribution of the frequency of individual clonotypes (including *TRAV/TRBV* to *TRAJ/TRBJ* pairing) was analyzed within unique sequences. Heat map representation of the frequencies of individual TRAV to TRAJ gene pairs and sequence overlap and treemaps representing CDR3 within the sample was produced using R software version 3.6.3 (2020-02-29). ImmunoSeqTM set of online tools was used to analyze the top 1000 most frequent clones, Shannon Entropy index of Diversity [H].

**Supplemental References.**


1. Kurgan G, Turk R, Li H, Roberts N, Rettig GR, Jacobi AM, Tso L, Sturgeon M, Mertens M, Noten R *et al*: CRISPAltRations: a validated cloud-based approach for interrogation of double-strand break repair mediated by CRISPR genome editing. *Mol Ther Methods Clin Dev* 2021, 21:478-491.
2. Noordermeer SM, Adam S, Setiaputra D, Barazas M, Pettitt SJ, Ling AK, Olivieri M, Alvarez-Quilon A, Moatti N, Zimmermann M *et al*: The shieldin complex mediates 53BP1-dependent DNA repair. *Nature* 2018, 560(7716):117-121.

3. Silva HM, Takenaka MC, Moraes-Vieira PM, Monteiro SM, Hernandez MO, Chaara W, Six A, Agena F, Sesterheim P, Barbe-Tuana FM *et al*: Preserving the B-cell compartment favors operational tolerance in human renal transplantation. *Mol Med* 2012, 18:733-743.
